# Physiologic and nanoscale distinctions define glutamatergic synapses in tonic vs phasic neurons

**DOI:** 10.1101/2022.12.21.521505

**Authors:** Kaikai He, Yifu Han, Xiling Li, Roberto X. Hernandez, Danielle V. Riboul, Touhid Feghhi, Karlis A. Justs, Olena Mahneva, Sarah Perry, Gregory T. Macleod, Dion Dickman

## Abstract

Neurons exhibit a striking degree of functional diversity, each one tuned to the needs of the circuitry in which it is embedded. A fundamental functional dichotomy occurs in activity patterns, with some neurons firing at a relatively constant “tonic” rate, while others fire in bursts - a “phasic” pattern. Synapses formed by tonic vs phasic neurons are also functionally differentiated, yet the bases of their distinctive properties remain enigmatic. A major challenge towards illuminating the synaptic differences between tonic and phasic neurons is the difficulty in isolating their physiological properties. At the *Drosophila* neuromuscular junction (NMJ), most muscle fibers are co-innervated by two motor neurons, the tonic “MN-Ib” and phasic “MN-Is”. Here, we employed selective expression of a newly developed botulinum neurotoxin (BoNT-C) transgene to silence tonic or phasic motor neurons. This approach revealed major differences in their neurotransmitter release properties, including probability, short-term plasticity, and vesicle pools. Furthermore, Ca^2+^ imaging demonstrated ~two-fold greater Ca^2+^ influx at phasic neuron release sites relative to tonic, along with enhanced synaptic vesicle coupling. Finally, confocal and super resolution imaging revealed that phasic neuron release sites are organized in a more compact arrangement, with enhanced stoichiometry of voltage-gated Ca^2+^ channels relative to other active zone scaffolds. These data suggest that distinctions in active zone nano-architecture and Ca^2+^ influx collaborate to differentially tune glutamate release at synapses of tonic vs phasic neuronal subtypes.

## INTRODUCTION

Most neurons are co-innervated by multiple inputs differing in neurotransmitter subtype, number of synaptic contacts, patterns of activity, and synaptic strength. This input-specific synaptic diversity, in turn, sculps patterns of activity in the target neuron to establish functional properties according to the demands of the neural circuitry in which it is embedded. These properties are not static, where synaptic inputs dynamically change throughout life and are crucial substrates for plasticity, modifying functional properties throughout development, maturation, experience, and aging. To fully understand input-specific synaptic properties, a major challenge is to isolate neurotransmission from individual inputs and characterize synaptic function in the absence of confounds emanating from other inputs.

The *Drosophila* neuromuscular junction (NMJ) is in principle a powerful model system to isolate and interrogate input-specific synaptic properties. At this final synapse in the motor circuit, most postsynaptic muscle fibers are co-innervated by two closely related motor neuron (MN) subtypes, MN-Is and MN-Ib, that differ in morphological and functional properties (Aponte-Santiago & Littleton, 2020; Johansen et al., 1989; Kurdyak et al., 1994; Lnenicka & Keshishian, 2000). Presynaptic terminals of phasic MN-Is are smaller in size, fire in bursts of activity, and typically innervate groups of muscle fibers to coordinate changes in locomotor activity (Choi et al., 2004; Schaefer et al., 2010). In contrast, tonic MN-Ib terminals are large, opposed by a complex subsynaptic reticulum, fire with relatively constant patterns of activity, and innervate only 1-2 muscle fibers to drive contraction (Aponte-Santiago & Littleton, 2020; Chouhan et al., 2010). Interestingly, neurotransmission at individual release sites of the “small” MN-Is input is strong, relative to those from the “big” MN-Ib input (Han et al., 2022; Lu et al., 2016; Newman et al., 2017). Yet it is unknown why these two developmentally-closely related glutamatergic synapses, sharing nearly identical molecular properties (Kohsaka et al., 2012), turn out to have such dramatic differences in synaptic strength.

Recently, new tools have been developed that enable electrophysiological isolation of MN-Is vs MN-Ib NMJs in *Drosophila*. In particular, GAL4 driver lines have been identified that enable selective expression in MN-Is or MN-Ib innervating muscle 6 (Aponte-Santiago et al., 2020; Carrillo-Reid et al., 2019; Pérez-Moreno & O’Kane, 2019). In addition, silencing of all neurotransmission from MN-Is or MN-Ib is now possible following the engineering of a botulinum neurotoxin (BoNT-C), which when selectively expressed in MN-Is or MN-Ib silences all neurotransmission without inducing heterosynaptic structural or functional plasticity at the convergent input (Han et al., 2022). While basic electrophysiological properties at MN-Is and MN-Ib NMJs were characterized with single action-potential evoked stimulation under low extracellular Ca^2+^ conditions using these tools (Han et al., 2022), the full range of synaptic physiological properties, and, importantly, why MN-Is forms stronger individual synapses relative to MN-Ib, has not been determined.

Here, we have isolated synaptic transmission at MN-Is vs MN-Ib NMJs using selective BoNT-C expression to characterize key physiological parameters including Ca^2+^ influx and cooperativity, vesicle pool size, and active zone nano-structure. We find that despite a reduced anatomical and functional release site number, active zones at strong MN-Is terminals experience enhanced Ca^2+^ influx, tighter vesicle coupling, and more compact active zone architecture with a higher ratio of CaV2 Ca^2+^ channels relative to weak MN-Ib terminals which together results in enhanced synaptic strength at these phasic inputs.

## RESULTS

### MN-Is functions as a high release probability synapse relative to MN-Ib

To understand why synaptic strength is stronger at MN-Is NMJs relative to those made by MN-Ib, we first undertook a systematic characterization of synaptic electrophysiology at isolated Is vs Ib NMJs. As described recently, selective expression of botulinum neurotoxin C (BoNT-C) disambiguates neurotransmission from MN-Is or MN-Ib, where evoked excitatory postsynaptic potential (EPSP) amplitude at low extracellular Ca^2+^ (0.4 mM) was ~2 fold higher at MN-Is compared to MN-Ib (Han et al., 2022). Synaptic strength can be described as a function of three variables: **n** (the number of release sites), **p** (the probability of synaptic vesicle fusion at a release site), and **q** (quantal size, the postsynaptic response to a single vesicle fusion). To assess these values, we used two-electrode voltage clamp (TEVC) electrophysiological recordings at isolated Is vs Ib neurons.

Consistent with our previous observations (Han et al., 2022), miniature excitatory postsynaptic current (mEPSC) amplitude (q) was ~50% larger at Is vs Ib (**Fig. 1D**), a difference for which mechanisms have already been established. Despite input-specific differences in the expression of glutamate receptor subtypes that should enhance postsynaptic sensitivity at Ib NMJs over Is (Han et al., 2022; Marrus et al., 2004), synaptic vesicles are larger at MN-Is terminals (Karunanithi et al., 2002), which leads to increased glutamate emission from each vesicle and drives the observed increase in quantal size. However, the 50% increase in q at Is NMJs is not sufficient, alone, to explain the enhanced synaptic strength of over 200%. Therefore, to better understand the higher neurotransmitter release at MN-Is terminals, we went on to assess synaptic strength across a range of extracellular Ca^2+^ concentrations (0.4 – 6.0 mM; **Fig. 1**).

**Figure 1:**
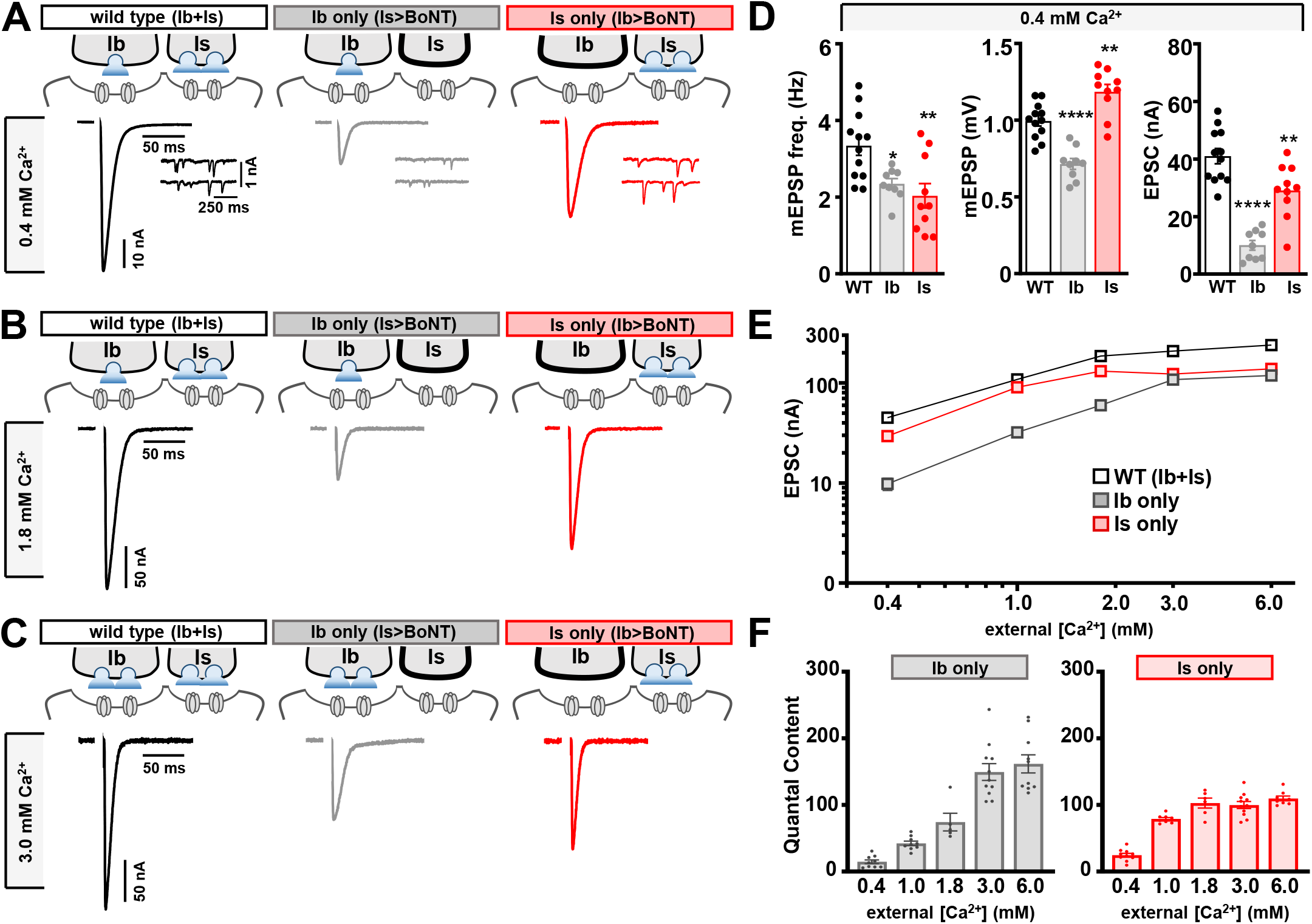
Presynaptic neurotransmitter release saturates at 1.8 mM Ca^2+^ at MN-Is terminals. **(A-C)** Schematic and representative EPSC and mEPSC traces of muscle 6 NMJ recordings from wild type (*w^1118^;* both Ib+Is stimulation), Ib only (Is>BoNT-C: *w;+;dHb9-GAL4/UAS-BoNT-C*), and Is only (Ib>BoNT-C: *w;+;GMR27E09-GAL4/UAS-BoNT-C*) at 0.4 mM (A), 1.8 mM (B), and 3.0 mM (C) extracellular Ca^2+^ concentrations. **(D)** Quantification of miniature frequency, amplitude, and evoked amplitude in wild type, Ib only, and Is only NMJs at 0.4 mM Ca^2+^. **(E)** Log-log plot of EPSC amplitude as a function of external [Ca^2+^] in the indicated genotypes. Note that while MN-Is saturates at [Ca^2+^]≥1.8 mM, transmission from MN-Ib continues to climb across the range of 0.4-6.0 mM Ca^2+^. **(F)** Plot of quantal content as a function of external [Ca^2+^] in Ib only and Is only NMJs. Error bars indicate ± SEM. ****p<0.0001; **p<0.01; *p<0.05. Additional statistical details are shown in Supplemental Table S1.

Our analyses revealed that at physiologic (1.8 mM Ca^2+^) and below, EPSC amplitudes from MN-Is NMJs were ~2x larger than MN-Ib (**Fig. 1A-E**). Above 1.8 mM Ca^2+^, presynaptic neurotransmitter release from MN-Is saturates, without a significant increase in EPSC amplitude or quantal content at 3-6 mM (**Fig. 1E,F**). In contrast, EPSC amplitude from MN-Ib NMJs increased across the conditions assayed, achieving parity with Is at 6 mM Ca^2+^ (**Fig. 1A-E**). While quantal content from MN-Is plateaued at 1.8 mM and above, MN-Ib released more quanta throughout the Ca^2+^ ranges tested (**Fig. 1F**). Thus, while synaptic strength at MN-Ib is inferior to MN-Is at physiological Ca^2+^ and below, tonic synapses have a much larger dynamic range and release does not saturate even at very high extracellular Ca^2+^ conditions.

The behavior of electrophysiologically isolated MN-Is vs MN-Ib NMJs suggest that probability of release (Pr) is relatively high at MN-Is terminals and low at MN-Ib. High vs low Pr synapses exhibit characteristic differences in short-term plasticity, with high Pr synapses depressing with repeated trains of stimulation and low Pr synapses facilitating (Karunanithi et al., 2002; Newman et al., 2017; Oleskevich et al., 2000) We tested these predictions by assessing paired-pulse plasticity (PPP) at low (0.4 mM) and high (6 mM) Ca^2+^ saline (**Fig. 2A,B**). As expected, PPP facilitated from MN-Ib NMJs and depressed at MN-Is (**Fig. 2A,B**). Furthermore, we measured synaptic augmentation with 30 stimuli at 60 Hz. EPSC amplitude steadily depressed at MN-Is NMJs in 3 mM Ca^2+^ conditions, while MN-Ib actually facilitated before slowly depressing (**Fig. 2C,D**). Given the differences in postsynaptic glutamate receptive fields at MN-Is vs -Ib NMJs (Han et al., 2022; Marrus et al., 2004), we sought to measure Pr using an approach that does not rely on postsynaptic sensitivity. Failure analysis is a method in which very low extracellular Ca^2+^ conditions are used so that about half of all stimulations leads to failures in evoked transmission; the relative frequency of failures can then be used to estimate Pr, with increased failures correlating with low Pr and lowered failures with high Pr (Huang & Stevens, 1997). Indeed, recording in 0.10 mM Ca^2+^, failure rates were 63% at MN-Is NMJs, but 95% at MN-Ib (**Fig. 3A,B**), confirming through both short-term plasticity and failure analysis that MN-Is indeed behaves like a high Pr synapse, while MN-Ib operates as a low Pr synapse.

**Figure 2:**
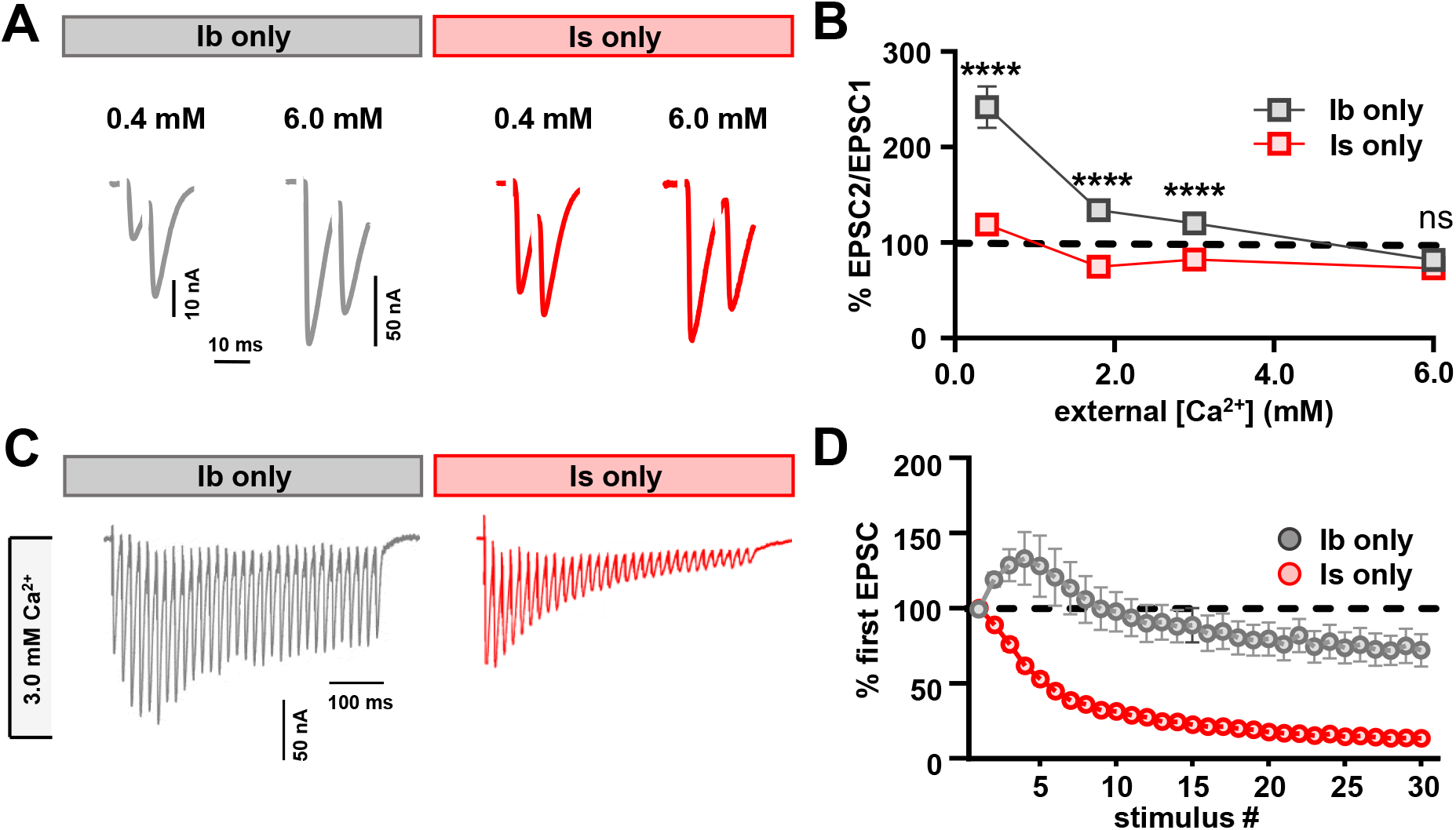
Short-term plasticity at isolated MN-Is and MN-Ib NMJs. **(A)** Representative paired-pulse EPSC traces of isolated Ib and Is NMJs recorded at 0.4 mM and 6.0 mM extracellular Ca^2+^ with a 16.7 msec interstimulus interval. **(B)** Paired-pulse ratios (EPSC2/EPSC1) plotted as a function of extracellular [Ca^2+^]. MN-Ib facilitates as lower Ca^2+^ levels, consistent with MN-Ib have a lower release probability relative to MN-Is (unpaired T-test). **(C)** Representative trains of 30 EPSCs stimulated at 60 Hz in isolated MN-Ib and MN-Is NMJs at 3.0 mM [Ca^2+^]. **(D)** EPSC amplitude for stimulation normalized to the amplitude of the initial EPSC. Note that while MN-Is depresses throughout the protocol, MN-Ib facilitates during the first 10 stimuli. Error bars indicate ± SEM. ****p<0.0001; (ns)=not significant. Additional statistical details are shown in Supplemental Table S1.

**Figure 3:**
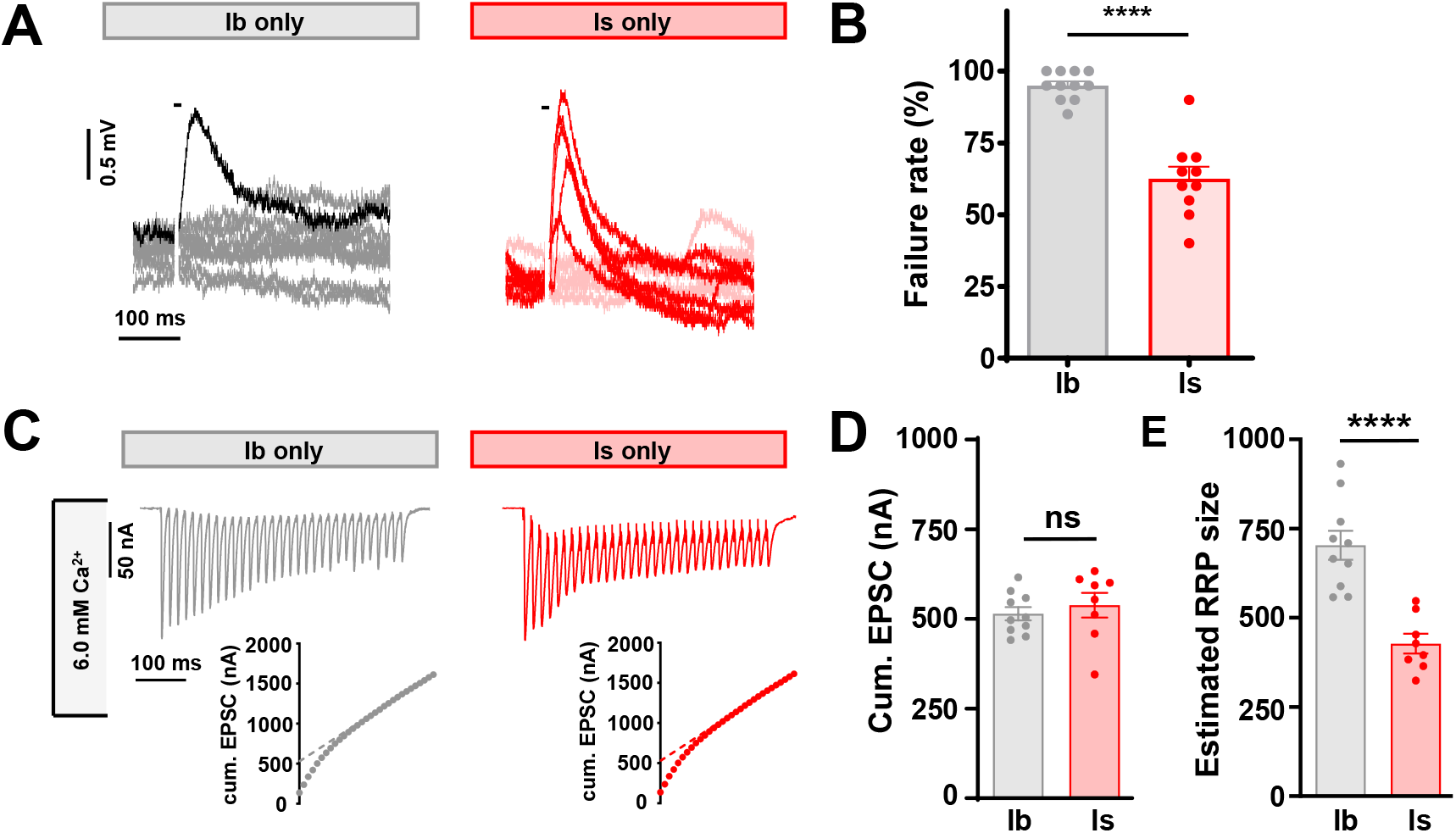
Failure analysis and determination of the readily releasable pool at isolated Is vs Ib synapses. **(A)** Representative traces of 10 stimulations of MN-Ib and MN-Is recorded at 0.10 mM Ca^2+^. The short line shows the time of stimulation, failures exhibit no responses. **(B)** Quantification of failure rate across 20 stimuli at 0.5 Hz in the indicated neurons. **(C)** Top: Representative traces of 30 EPSCs from MN-Ib and MN-Is at 60 Hz stimulation in 6 mM extracellular Ca^2+^. **Bottom:** Averaged cumulative EPSC amplitudes. A line fit to the 18-30^th^ stimuli was back extrapolated to time 0. **(E)** Quantification of the average cumulative EPSC values and estimated readily releasable pool (RRP) sizes at MN-Ib and MN-Is. Error bars indicate ± SEM. ****p<0.0001; (ns)=not significant. Additional statistical details are shown in Supplemental table S1.

Finally, we sought to define the relative readily-releasable vesicle pool (RRP) at MN-Is vs -Ib. The RRP is functionally defined as the subset of vesicles at presynaptic boutons that are released upon a rapid burst of activity (Kaeser & Regehr, 2017; Schneggenburger et al., 1999), and can be estimated by using 30 stimuli of high frequency (60 Hz) stimulation at elevated extracellular Ca^2+^ (6 mM) (**Fig. 3C**). The cumulative EPSC is plotted, where back-extrapolation with the linear phase can be used to estimate the RRP size (Goel, Li, et al., 2019). Using this approach, we determined that the strong MN-Is NMJ has ~half the RRP relative to MN-Ib (**Fig. 3C-E**). Together, these data establish two major electrophysiological distinctions between MN-Is vs -Ib synapses: First, the higher quantal size and Pr at MN-Is together are the major contributors to the enhanced synaptic strength observed at phasic inputs. Second, the weak MN-Ib is capable of a much larger dynamic range of quantal content, where additional neurotransmitter release is facilitated during high activity using a larger RRP to mobilize additional vesicles for fusion.

### Release site number is enhanced at the weak MN-Ib

Next, we sought to assess the number of synaptic vesicle release sites, “n”, at MN-Is vs -Ib terminals. Release sites can be anatomically or functionally defined. One anatomical measure of release sites corresponds to active zones (AZs), areas of electron dense presynaptic scaffolding, where the major “T-bar” scaffold Bruchpilot (BRP) can be used to label individual AZs (Kittel et al., 2006; Wagh et al., 2006). Using confocal imaging, we immunostained BRP at MN-Is and -Ib NMJs innervating muscle 6 (segment A3), where all of the electrophysiology in this study was conducted, and quantified total BRP puncta number per NMJ to determine anatomical release site number. We found that while the weak MN-Ib had ~200 BRP puncta/M6 NMJ, the strong MN-Is had less than 100 (**Fig. 4A,B**). Thus, although more neurotransmitter is released from strong MN-Is terminals, there are less than half the number of anatomical release sites.

**Figure 4:**
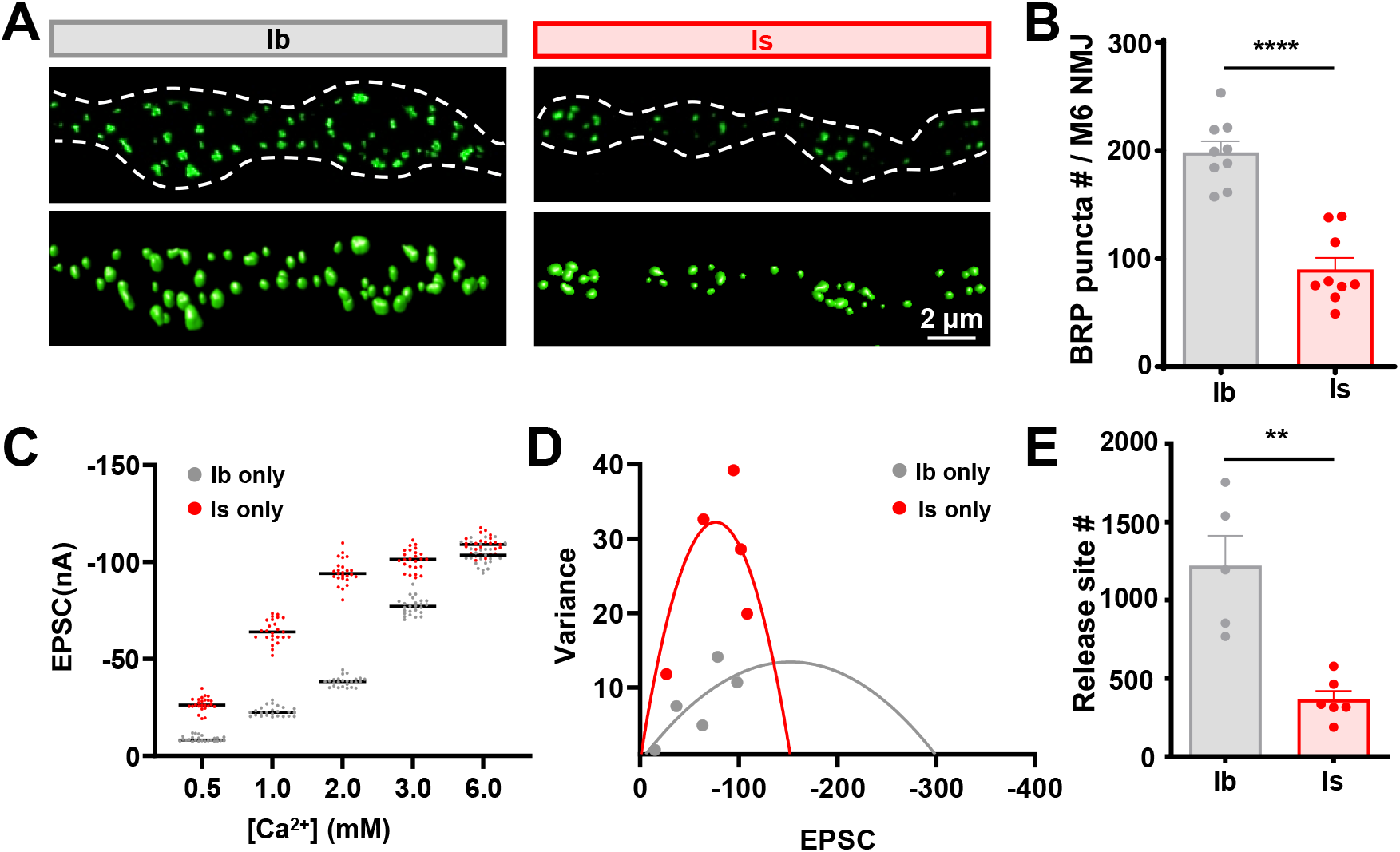
Determination of anatomic and functional release sites at disambiguated MN-Ib and MN-Is terminals. **A)** Representative confocal images of MN-Ib and MN-Is boutons immunostained with antibodies recognizing the active zone scaffold BRP, with dash lines indicating the neuronal membrane stained with anti-HRP. Below: Surface renderings from confocal images of individual BRP puncta. **(B)** Quantification of BRP puncta number per entire muscle 6 from MN-Ib and MN-Is NMJs. A twofold increase in BRP puncta number is observed at MN-Ib compared to MN-Is. **(C)** Scatter plot EPSC amplitude distribution from MN-Ib and MN-Is in the indicated extracellular [Ca^2+^]. **(D)** Variance-mean plots for the data shown in (C). Variance was plotted against the mean amplitude of 10 EPSCs recorded at 0.2 Hz from the five Ca^2+^ concentrations detailed in (C). **(E)** Estimated number of functional release sites based on the mean-variance plots in (D), with a >threefold increase at MN-Ib relative to MN-Is. Error bars indicate ± SEM. ****p<0.0001; **p<0.01. Additional statistical details are shown in Supplemental Table S1.

Although there is a reduced anatomical AZ number at MN-Is, “n” more specifically refers to the number of synaptic vesicle (SV) fusion sites embedded within all AZs, likely defined by Unc13 (Jahn & Fasshauer, 2012; Reddy-Alla et al., 2017; Sakamoto et al., 2018) Recent studies have shown that a significant fraction of anatomical release sites are functionally silent for evoked fusion at the fly NMJ, and that Pr varies widely across AZs (Akbergenova et al., 2018; Gratz et al., 2019; Melom et al., 2013; Newman et al., 2022). Mean-variance analysis is a mathematical approach that uses electrophysiological recordings over a range of extracellular Ca^2+^ conditions to measure the variance of EPSC amplitudes (Clements & Silver, 2000; Li et al., 2018; Sola et al., 2004). The probabilistic nature of synaptic vesicle fusion is used to extract the number of functional release sites from the EPSC variance (Clements, 2003). Using this approach, we determined that MN-Is had about one third the number of functional release sites compared to MN-Ib, ~400 vs ~1200 (**Fig. 4C-E**). Thus, both anatomical and functional release site numbers favor enhanced release at MN-Ib over MN-Is. Together, these data demonstrate that despite MN-Ib having increased number of release sites, both quantal size and release probability are higher at MN-Is, contributing to the observed increase in synaptic strength from phasic inputs.

### Ratiometric Ca^2+^ imaging reveals functional differentiation between terminal subtypes

Next, we sought to understand the mechanisms responsible for the enhanced release probability exhibited by MN-Is discussed above. A major determinant of Pr is the abundance and dynamics of Ca^2+^ entry into presynaptic terminals after an action potential. Here, we used quantitative ratiometric Ca^2+^ imaging techniques with chemical dyes forward-filled into Ib and Is terminals to determine the extent to which differences in Ca^2+^ influx at AZs at MN-Is might explain the higher Pr at phasic release sites (**Fig. 5**).

**Figure 5:**
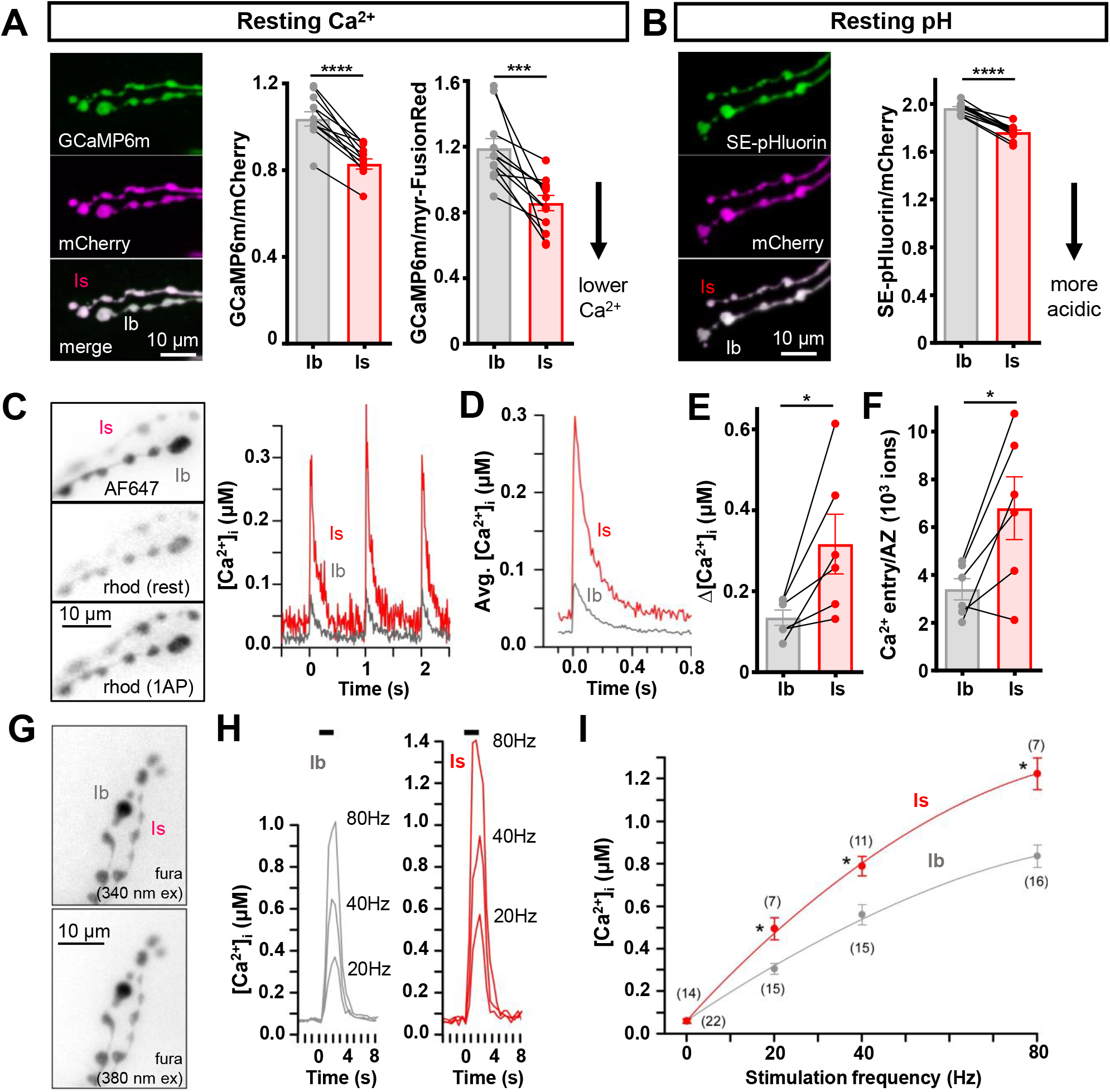
Single-action-potential-evoked Ca^2+^ transients are two-fold higher at MN-Is release sites. **(A,B)** Resting [Ca^2+^] (A) and pH (B) estimated in intact NMJ boutons using two previously established genetically encoded reporters (Ca^2+^: Gerry: (Ugur et al., 2017) and myrFusionRed-GCaMP6m: (Stawarski et al., 2020); pH: pHerry: (Rossano et al., 2017). **(A)** Images of the “Gerry” fusion protein, consisting of GCaMP6m C-terminally linked to mCherry, expressed in both terminal types using the nSyb-Gal4 driver (*nSyb-GAL4>GCaMP6m-mCherry*). Middle: Plots of the average GCaMP6m:mCherry fluorescence ratio measured in the cytosol of terminal pairs. Right: Plots of the average GCaMP6m to FusionRed fluorescence ratio in the cytosol of terminal pairs. The C-terminus of myristoylated FusionRed was fused to the N-terminus of GCaMP6m (*OK6-GAL4>myrFusionRed-GCaMP6m*). Terminal images not shown. Is boutons have an apparent lower resting [Ca^2+^] compared to Ib (Middle: N=11; P < 0.001; Left: N=12; P < 0.001; paired Student’s T-tests). **(B)** Images of the “pHerry” fusion protein, consisting of SE-pHluorin C-terminally linked to mCherry, expressed in both terminal types using the *nSyb-Gal4* driver (*nSyb-GAL4>SE-pHluorin-mCherrÿ*). Right: Plots of the average SE-pHerry:mCherry fluorescence ratio measured in the cytosol of terminal pairs. Is terminals are slightly more acidic relative to Ib (N=12; P < 0.001, paired Student’s T-test). **(C-F)** Average Ca^2+^ entry per AZ per AP estimated using forward-filled Ca^2+^-sensitive dye rhod and Ca^2+^-insensitive dye AF647. **(C)** Inverted grayscale images of AF647 or rhod fluorescence from a pair of Ib and Is bouton terminals. Right: Traces revealing the transient increase in free Ca^2+^ concentration ([Ca^2+^]) at Is vs Ib boutons in response to the first 3 of 10 nerve impulses delivered at 1 Hz. [Ca^2+^] was estimated from an *in situ* calibration of rhod fluorescence relative to Ca^2+^-insensitive AF647 (see Methods). **(D)** Traces representing the numerical average of [Ca^2+^] during 10 APs at each terminal shown in C. **(E)** Plots of the average maximum change in [Ca^2+^] in response to an AP (Δ[Ca^2+^]_AP_) at terminal pairs in 6 separate preparations. Average Δ[Ca^2+^]_AP_ was higher for Is terminals (N=6; P = 0.034, paired Student’s T-test). **(F)** Plots of the average Ca^2+^ entry (number of ions) per AZ per AP (see Methods) (P<0.024, paired Student’s T-test). (**G-I**) Peak [Ca^2+^] achieved during nerve stimulus trains estimated using forward-filled Ca^2+^-sensitive dye fura. (**G**) Inverted grayscale images of fura fluorescence from terminals on muscle fiber #6. Each image is the average of 10 frames collected at 2 fps, prior to nerve stimulation, using either 340 nm or 380 nm excitation, while collecting emission at 510 nm. (**H**) Traces revealing the increase in [Ca^2+^] in response to a series of 2 second trains with different frequencies (20, 40 and 80 Hz). [Ca^2+^] was estimated from an *in situ* calibration of fura fluorescence ratios (see Methods). (**I**) Plots of the average maximum change in [Ca^2+^] in response to trains. Numbers in parentheses represent the number of separate preparations (two-way ANOVA, p<0.05 overall, Holm-Sidak post hoc test). All experiments performed on muscle 6. Measurements in A-B made in 0.1 mM Ca^2+^, 15 mM Mg^2+^ HL6, while measurements in C-I made in 2 mM Ca^2+^, 15 mM Mg^2+^ HL6. Error bars indicate ± SEM. ****p<0.0001; ***p<0.001; *p<0.05. Additional statistical details are shown in Supplemental Table S1.

Previous studies have estimated a ~3-fold higher average probability of release from AZs in type-Is vs -Ib (p=0.3 vs 0.1; (Akbergenova et al., 2018; Lu et al., 2016; Newman et al., 2017). This might be explained by differences in the amount of Ca^2+^ that enters per AZ during an action potential, or by differences in the “global” concentration of free Ca^2+^ inside the terminal ([Ca^2+^]_i_) at rest. Elevated global [Ca^2+^]_i_ is associated with an accelerated recruitment of release-ready SVs (Neher & Sakaba, 2008), and so we determined whether type-Is terminals maintain a higher [Ca^2+^]_i_ at rest that might enhance SV docking and priming. Unfortunately, our findings were inconclusive due to limitations in the properties of both chemical and genetically-encoded Ca^2+^-reporters, and the methods of their application. When the ratio of Ca^2+^-sensitive rhod to Ca^2+^-insensitive AF647 was examined in type-Is terminals, a higher resting [Ca^2+^]_i_ was observed (Is: 37.2 ± 9.3 nM; Ib: 23.9 ± 6.4; SEM, P=0.023; Student’s T-test), consistent with a larger data set previously published using OGB1 in ratio to AF568 (Han et al., 2020). Curiously, such a difference was not supported by a similarly robust fura data set (Fig. 5I; P=0.99, two-way ANOVA). To guard against the possibility that the differences in resting [Ca^2+^]_i_ shown by both rhod/AF647 and OGB1/AF568 reflect differential tolerance to the axotomy that takes place up to 6 hours prior to measurements with forward-filled chemical Ca^2+^ reporters, we also used genetically-encoded Ca^2+^ reporters with axons intact. Contrary to rhod- and OGB1-based measurements, the fluorescence ratio of Ca^2+^-sensitive GCaMP6m to either Ca^2+^-insensitive mCherry or myristoylated-FusionRed indicated a higher resting [Ca^2+^]_i_ in type-Ib terminals (**Fig. 5A**). However, we believe this difference to be the result of greater quenching of GCaMP6m fluorescence in type-Is terminals as Ca^2+^-bound GCaMP6m is highly-sensitive to pH change near neutral (Barnett et al., 2017) and we discovered type-Is terminals to be more acidic using pH-sensitive SE-pHluorin expressed freely in the cytosol in tandem with mCherry (**Fig. 5B**). This pH difference might reconcile the fura data with the rhod and OGB-1 data above, as fura is more sensitive to pH change near neutral than either rhod or OGB-1 (Geisow & Evans, 1984; Grynkiewicz et al., 1985; ThermoFisher, 2022) in which case [Ca^2+^]_i_ may be underestimated in fura-filled type-Is terminals. This difference in pH between terminals is a tantalizing finding, as intracellular acidification is capable of either facilitating or depressing Ca^2+^ current in mammalian neurons (Saegusa et al., 2011; Tombaugh & Somjen, 1997) and although its effect on Ca_v_2 channel Cac in these terminals is not known, it is a candidate mechanism for the greater Ca^2+^ influx through type-Is AZs.

Using the techniques described by Lu and others (2016) we made estimates of Ca^2+^ entry per AZ at the different terminal types. Initially, we quantified changes in fluorescence (ΔF) of an intracellular Ca^2+^-sensitive dye (rhod) in response to single APs, and then calibrated rhod ΔF relative to changes in [Ca^2+^]_i_ (Δ[Ca^2+^]_i_; **Fig. 5C-E**). Through reference to the amount of dye loaded, endogenous Ca^2+^ buffering capacity, bouton volume and AZ numbers quantified by Lu and others (2016), we estimated that twice as many Ca^2+^ ions enter through AZs on type-Is terminals (Is: [6.8 ± 1.3] x 10^3^; Ib: [3.4 ± 0.4] x 10^3^; SEM, P = 0.024; paired Student’s T-test; **Fig. 5F**). This result is consistent with the findings of Lu and others (2016). Due to the power dependence of transmitter release on Ca^2+^ entry at this NMJ and other synapses (Dodge & Rahamimoff, 1967; Jan & Jan, 1976) the two-fold greater Ca^2+^ entry at type-Is AZs is sufficient to explain the three-fold greater release probability at type-Is AZs. Using the Ca^2+^-sensitive dye fura, as described by Chouhan and others (Chouhan et al., 2010), we quantified the level to which the intracellular Ca^2+^ concentration [Ca^2+^]_i_ rises during short trains of APs (**Fig. 5G-I**). Here also, consistent with greater Ca^2+^ entry per AZ at type-Is terminals, and consistent with previous data collected from type-Ib/Is pairs on muscle #6 using OGB-1 (Han et al., 2020; He et al., 2009), type-Is terminals accumulate Ca^2+^ to higher levels during AP trains (**Fig. 5I**; *, P<0.001, by two-way ANOVA). Together, this quantitative analysis of Ca^2+^ dynamics demonstrates that AZs at MN-Is terminals experience ~2x increased Ca^2+^ influx per AZ relative to those at MN-Ib, sufficient to explain the increased Pr observed at phasic terminals.

### Active zone nano-architecture is compressed at MN-Is active zones

Given the differences in Ca^2+^ influx at tonic vs phasic release sites, we next assessed AZ structure to understand the features that contribute to the differential release characteristics at these distinct terminals. First, we examined AZ components by immunostaining and confocal microscopy. In particular, we labeled the T-bar scaffold BRP, the core AZ scaffold Rim binding protein (RBP), and the Cav2 voltage-gated Ca^2+^ channel Cac. We considered that perhaps an increased abundance of Cac at MN-Is AZs may explain the enhanced Ca^2+^ influx at phasic AZs. However, we found no significant difference in fluorescence intensity of Cac puncta endogenously labelled with GFP at MN-Is vs -Ib AZs (**Fig. 6A,C**). Despite the enhanced release at MN-Is AZs, we unexpectedly found that the florescence intensity of both BRP and RBP were diminished at MN-Is compared to -Ib (**Fig. 6A,C**), an intriguing contrast to what was observed for Cac, which was similar at AZs of both MNs. We also noted a small but significant increase in the density of AZs, labeled by BRP puncta, at MN-Is vs -Ib terminals (**Fig. 6A,B**), as observed previously (He et al., 2009). These characteristic properties of MN-Is AZs, with increased density and reduced intensity, could emerge as a result of the distinct tonic vs phasic activity patterns between Is vs Ib (Aponte-Santiago & Littleton, 2020; Choi et al., 2004; Chouhan et al., 2010; Kurdyak et al., 1994) We therefore perturbed activity patterns by chronic expression of the inwardly rectifying potassium channel *Kir2.1* in both MN-Is and -Ib (**Fig. S1A**). However, we observed the same characteristic differences in AZ density and BRP/RBP intensity at MN-Is vs - Ib despite chronic hypo-excitability induced by *Kir2.1* expression (**Fig. S1A-C**). Together, these results demonstrate two important points. First, tonic vs phasic activity patterns are not required to instruct the distinguishing characteristics of AZs at MN-Is vs -Ib, indicating that they are generated through activity-independent processes. Second, strong MN-Is terminals, counterintuitively, have reduced AZ number per NMJ (**Fig. 4A,B**) and diminished abundance of AZ scaffolds but retain similar abundance of CaV2 voltage-gated Ca^2+^ channels relative to MN-Ib release sites.

**Figure 6:**
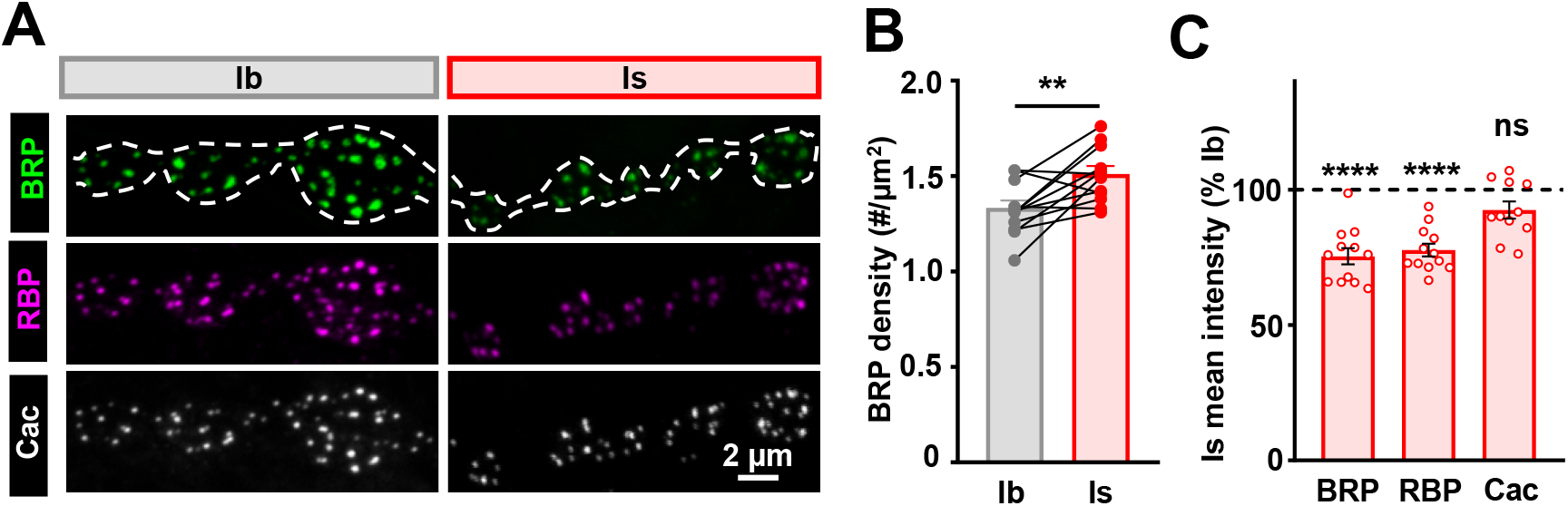
Active zone components are dimmer at MN-Is terminals compared to MN-Ib. **(A)** Representative confocal images of muscle 6 NMJs immunostained with antibodies against the active zone components BRP, RBP, and Cac (GFP; endogenously tagged Cac^sfGFP^; *cac^sfGFP^*^-N^; +; +) at MN-Ib and MN-Is. The dashed line represents the neuronal membrane indicated by the anti-HRP signal. **(B)** Quantification of BRP puncta density in paired terminals of the same muscle. BRP puncta were denser at MN-Is (paired Student’s T-test). **(C)** Immunofluorescence intensity of the indicated antibodies at MN-Is normalized as a percentage of that at MN-Ib. Note that while the intensity of BRP and RBP are significantly lower at MN-Is compared to MN-Ib, no significance difference is observed for Cac (paired Student’s T-test). Error bars indicate ± SEM. ****p<0.0001; **p<0.01; (ns)=not significant. Additional statistical details are shown in Supplemental Table S1.

The intriguing difference between Cac and the scaffolds RBP and BRP at MN-Is vs -Ib AZs suggested the existence of nano-architectural distinctions. Therefore, we next examined AZ nanostructure at Is vs Ib AZs using super resolution STimulated Emission Depletion (STED) microscopy. First, we assessed the dimensions of BRP, which at Ib terminals exhibit a characteristic “doughnut” organization with a ring surrounding a central hole, which is where Cac channels are clustered (Fouquet et al., 2009). In contrast to BRP rings at Ib, BRP was also organized in a annulus-like structure at MN-Is AZs, but with more of a triangular appearance and a marked reduction in size, exhibiting about half the area of Ib BRP rings (**Fig. 7A,B**). Within individual BRP rings, “nanomodules” of local intensities have been described (Böhme et al., 2019; Goel, Bergeron, et al., 2019; Muttathukunnel et al., 2022), with ~4 nanomodules per Ib BRP ring, consistent with our findings (**Fig. 7A,B**). Not only were BRP rings at Is reduced in area, but nanomodules were reduced to ~3 per BRP ring. Next, we probed BRP, RBP, and Cac dimensions and inter-relationships at individual Ib vs Is AZs. Consistent with previous Ib AZ studies (Fouquet et al., 2009; Liu et al., 2011), we found the same relationships held at Is AZs, with a central punctum of Cac surrounded by RBP scaffolding, all encompassed by a BRP ring at the periphery (**Fig. 7C,E**). However, while the same relative relationships held at both Ib/Is AZs, the dimensions were more compact at Is AZs, with parameters at both planar (top) and vertical (side) views ~10% smaller in both lateral and Z-dimensions (**Fig. 7C,E,F**). Finally, we considered the finding that while Cac intensity was similar at Ib vs Is AZs, both BRP and RBP intensity was reduced at Is (**Fig. 6C**). Using STED microscopy, we calculated the ratio of the area of RBP to BRP, and determined that these ratios were the same at Ib vs Is AZs (**Fig. 7D**). However, the area ratio of Cac to BRP was ~2-fold enhanced at Is AZs relative to Ib (**Fig. 7D**). These data indicate an enhanced stoichiometry of the voltage-gated Ca^2+^ channel Cac relative to the central AZ scaffolds and an overall compaction of AZ nano-structure at the strong MN-Is in relation to the weaker MN-Ib.

**Figure 7:**
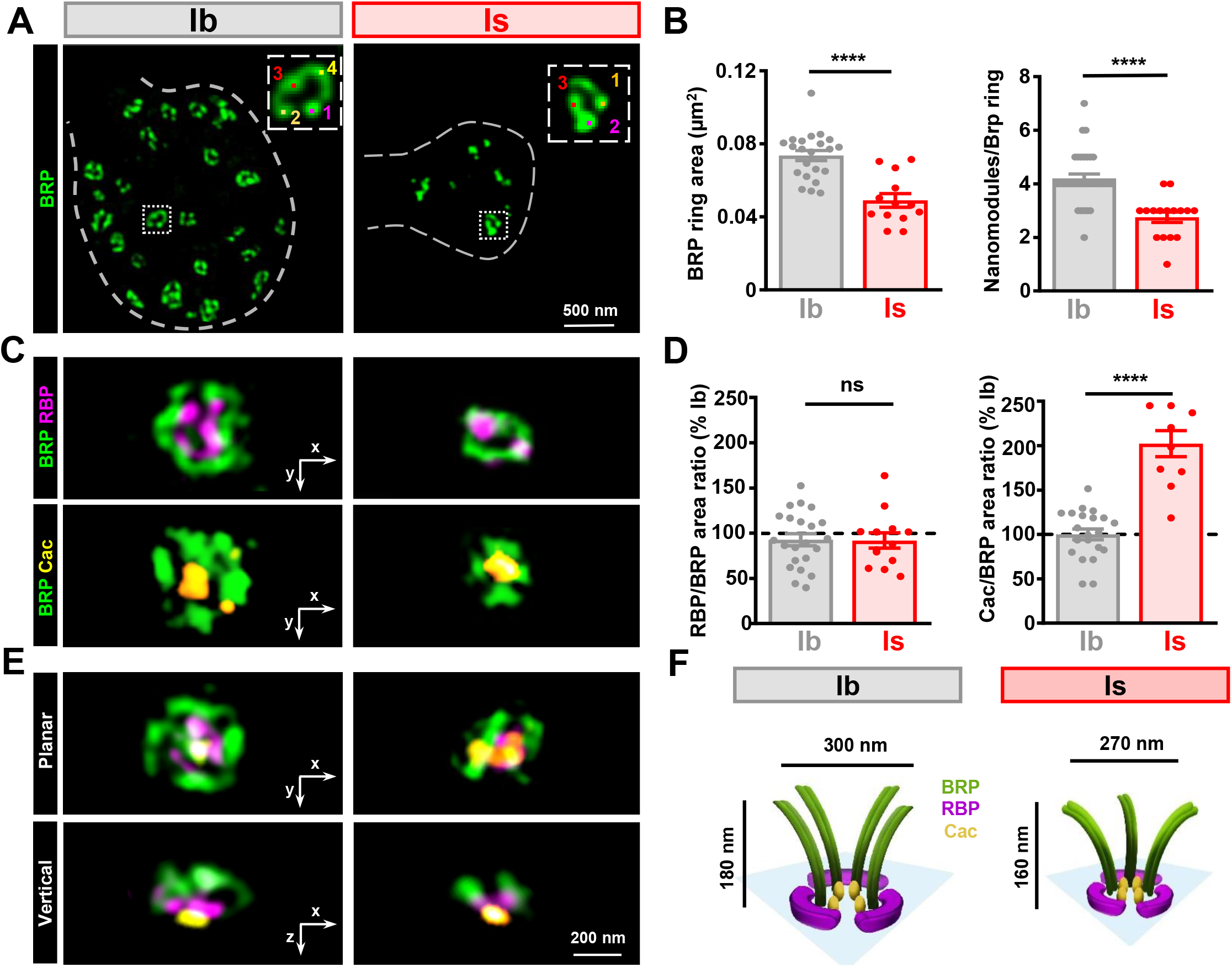
STED imaging reveals enhanced Ca_V_2 Ca^2+^ channel stoichiometry at MN-Is active zones. **(A)** Representative STED images of single NMJ boutons immunostained with anti-BRP at MN-Ib and MN-Is. The BRP signal appears as rings and has distinguishable “nanomodules” of local intensity maxima (Bohme et al., 2019). **(B)** Quantification of BRP ring area and number of nanomodules per BRP ring. MN-Is exhibits significantly smaller BRP rings and a concomitant reduction in BRP nanomodules. **(C)** Representative STED images of single active zones illustrating BRP and RBP (top) or BRP and Cac (bottom) at MN-Ib and MN-Is terminals. **(D)** Quantification of the RBP/BRP area ratio and Cac/BRP area ratio normalized to the average value of MN-Ib. MN-Is showed a twofold increase in the Cac/BRP area ratio compared to MN-Ib, while no difference was observed for RBP/BRP. **(E)** Representative STED images of single active zones triple stained with anti-BRP, -RBP, and -Cac (GFP) at planar (top) and vertical (bottom) orientations at MN-Ib and MN-Is terminals. **(F)** Schematics illustrating the spatial relationships between BRP, RBP, and Cac at active zones of MN-Ib and MN-Is. The diameter and height data are the average values of STED images at planar or vertical orientations. Error bars indicate ± SEM. ****p<0.0001; (ns)=not significant. Additional statistical details are shown in Supplemental Table S1.

### Vesicle-Ca^2+^ channel coupling is tighter at strong MN-Is release sites

The two-fold difference in presynaptic Ca^2+^ influx per AZ at MN-Is vs -Ib might be more than sufficient to explain the observed differences in release, as neurotransmitter release is a Ca^2+^-dependent cooperative process. At the fly larval NMJ, release from MN-Ib and -Is combined has been variously described as having a 3^rd^ or 4^th^ power dependence on extracellular Ca^2+^ (Bao et al., 2005; Jan & Jan, 1976; Paddock et al., 2008; Schneggenburger & Neher, 2005; Stewart et al., 2000). However, this power relationship only holds at lower values of extracellular Ca^2+^. To make the argument that MN-Ib AZs are equivalent to MN-Is AZs with a reduced Ca^2+^ influx, we would predict that doubling extracellular Ca^2+^, and Ca^2+^ entry, at a MN-Ib AZ would lead to a MN-Is AZ level of release, i.e. a 3-fold increase in release, and this seems unlikely on the basis of our empirical data in **Fig. 1F**. Given that we found evidence for similar abundance of Cac per AZ at Is vs Ib (**Fig. 6A,C**) and a more compressed nanostructure (**Fig. 7**), we hypothesized that synaptic vesicle coupling to Ca^2+^ channels may provide an additional layer of regulation to tune release properties and help to explain the differences in synaptic strength between MN-Ib vs -Is, as observed in other systems (Millar et al., 2005). If this were the case, we surmised that this coupling, and overall synaptic release properties, would be calibrated to the *in vivo* ionic conditions, namely Ca^2+^ and Mg^2+^ levels. Ionic concentrations of *Drosophila* hemolymph were determined decades ago using ion-sensitive electrodes following “bled” larvae (Stewart et al., 1994), which determined a [Ca^2+^] of 1.8 mM. However, Mg^2+^-sensitive electrodes were not available, and the concentration of 20 mM Mg^2+^ was chosen based largely on literature available at the time.

We therefore first decided to revisit these values using an independent approach. We engineered a low-affinity genetically-encoded Ca^2+^ indicator, CEPIA1er (Suzuki et al., 2014), capable of accurately reporting the relatively high extracellular Ca^2+^ levels in *Drosophila* larvae *in vivo*. We included a signal sequence to traffic CEPIA1er to the outer leaflet of the plasma membrane by fusing it to the human platelet-derived growth factor receptor (hPDGFR) transmembrane domain (TM) (**Fig. 8A**). For ratiometric imaging, we also included the Ca^2+^-insensitive fluorophore, TagRFP fused to hPDGFR-TM following a P2A sequence to ensure an equimolar expression ratio from the same transcript. We expressed this transgene in larval motor neurons and calibrated the CEPIA1er/RFP ratio through the cuticle of filet-dissected preparations across known Ca^2+^ concentrations in 15 mM Mg^2+^ (**Fig. 8B**). We then imaged CEPIA1er/RFP through the larvae cuticle *in vivo*, determining a [Ca^2+^] of 2.2 mM (**Fig. 8B,C**). However, given that the *in vivo* Mg^2+^ concentration remains unclear and that 10 mM Mg^2+^ is the standard saline concentration used for electrophysiology in the lab (Dickman et al., 2005; Goel, Li, et al., 2019; Kiragasi et al., 2017), we determined that 2.2 mM Ca^2+^/15 mM Mg^2+^ is equivalent to 1.8 mM Ca^2+^/10 mM Mg^2+^(**Table S1**). Thus, we used 1.8 mM Ca^2+^/10 mM Mg^2+^ conditions to probe vesicle coupling in our final set of experiments.

**Figure 8:**
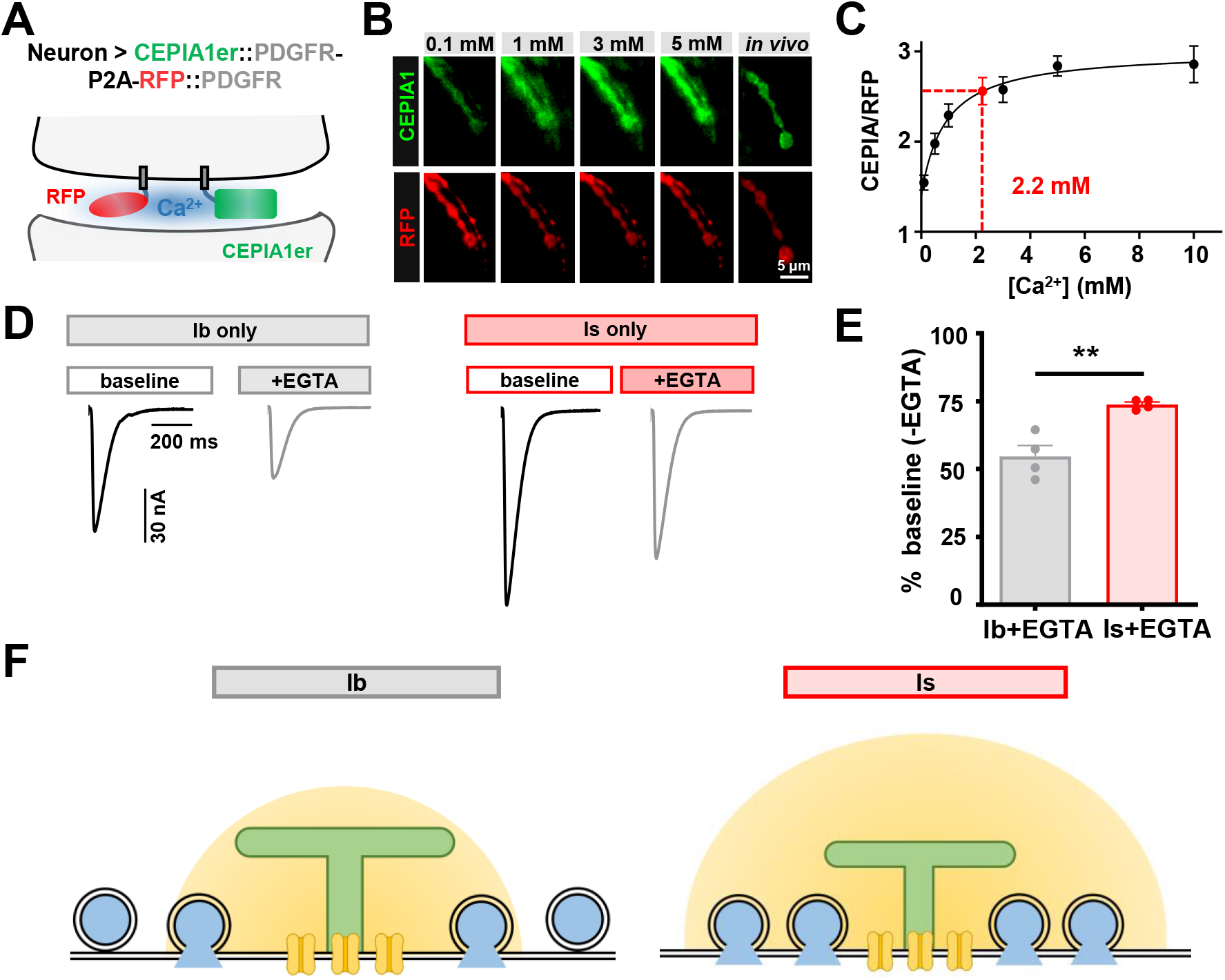
Synaptic vesicles are more tightly coupled at MN-Is release sites. **(A)** Schematic of high affinity Ca^2+^ indicator CEPIAer fused to the hPDGFR-TM (*nSyb-GAL4>CEPIA1er-hPDGFR-TM-P2A-RFP-hPDGFR-TM*), expressed in motor neurons in equimolar proportion with RFP also fused to the hPDGFR-TM. Note that both RFP and CEPIA1er are expressed in the extracellular space. **(B)** Representative live confocal images of NMJs expressing CEPIA1er and RFP at the indicated extracellular [Ca^2+^] in semi-dissected preparations for calibration, and *in vivo* from intact larvae. **(C)** Michaelis-Menten fitting of the CEPIA/RFP fluorescence intensity ratio plotted against [Ca^2+^]. The *in vivo* Ca^2+^ concentration (red dot) was determined to be 2.2 mM by locating the CEPIA/RFP value in the curve. **(D)** Representative EPSC traces at 1.8 mM Ca^2+^ illustrating the effect of acute application of EGTA-AM on isolated Ib or Is muscle 6 NMJs. **(E)** EPSC amplitude after EGTA treatment normalized to baseline (-EGTA). Synaptic release from MN-Ib is more sensitive to EGTA than release from MN-Is. **(F)** Schematics illustrating the larger EGTA-sensitive synaptic vesicle pool at Ib terminals. Error bars indicate ± SEM. **p<0.01. Additional statistical details are shown in Supplemental Table S1.

Synaptic vesicle-Ca^2+^ channel coupling typically refers to the physical distance between the Ca^2+^-trigger for release associated with synaptic vesicles and the source of Ca^2+^ (Dittman & Ryan, 2019; Neher & Sakaba, 2008; Wadel et al., 2007). “Tightly” coupled release sites have vesicle pools close to the Ca^2+^ sources and release more rapidly, while “loosely” coupled sites are further away. Relative vesicle coupling can be probed using the Ca^2+^ chelator EGTA-AM, which can be trapped in presynaptic terminals to compete with the Ca^2+^ trigger for access to Ca^2+^ (Genç et al., 2017; Hagler & Goda, 2001). Using this approach, presynaptic vesicle release is more diminished at AZs with looser coupling. Indeed, we observed that weak MN-Ib NMJs were more impacted by EGTA treatment compared to those of MN-Is, where equivalent concentrations of EGTA reduced EPSC amplitude by ~50% at Ib but only ~25% at Is (**Fig. 8D,E**). Hence, synaptic vesicles are more tightly coupled at the strong MN-Is, which likely works with enhanced Ca^2+^ influx to tune release properties despite reduced overall number of anatomic and functional release site number, but saturates above physiological Ca^2+^ levels. In contrast, less release is observed at MN-Ib at physiological Ca^2+^ and below, but more release continues above this threshold as more Ca^2+^ influx engages more loosely coupled vesicles.

## DISCUSSION

By disentangling the electrophysiological properties of synapses made by tonic and phasic MNs at the *Drosophila* NMJ, we have revealed their distinct functional and structural properties. At physiologic conditions, phasic MN-Is operates at its maximal capacity, functioning as a “high Pr” synapse with enhanced Ca^2+^ influx and depressing with repeated trains of stimulation. In contrast, the tonic MN-Ib possesses a much larger dynamic range, exhibiting relatively low release probability but facilitating and “ramping up” as firing bouts continue. Form follows function, and their AZ topologies are similarly specialized, with compact AZs organized at phasic terminals biased to release tightly coupled vesicles and larger tonic AZs engaging more loosely coupled vesicles. These neuronal subtypes were molded through evolution to meet the specific demands of the neural circuitry in which they are embedded to coordinate larval locomotor behavior.

### Presynaptic release properties of MN-Is vs -Ib

Two physiologic properties render phasic MN-Is the “stronger” synapse in terms of EPSC amplitude evoked with single action potentials. First, the larger volume of individual synaptic vesicles at Is terminals generates a quantal size (**q**) ~50% larger than Ib (Han et al., 2022; Karunanithi et al., 2002; Lu et al., 2016) Secondly, increased release probability (**p**), driven by enhanced Ca^2+^ influx per AZ and more tightly coupled vesicles, also strengthens transmission at MN-Is over -Ib. These two properties combine to overcome that property which favors synaptic strength at tonic MN-Ib – an enhanced numbers of release sites (**n**). Low release probability, enhanced release site number and great number of vesicles available for release (Justs et al., 2022)all favor facilitation at tonic MN-Ib inputs vs depression at phasic MN-Is inputs, consistent with previous work (Justs et al., 2022; Kurdyak et al., 1994; Lu et al., 2016; Newman et al., 2017) The distinctions in tonic vs phasic synaptic transmission is likely related to the functions of the MN subtypes in orchestrating larval locomotion, where individual tonic MN-Ib inputs innervate specific muscle fibers to drive contraction, while phasic MN-Is innervates groups of muscles to coordinate changes in direction (Heckscher et al., 2012; Schaefer et al., 2010) Hence, the physiological signatures of tonic and phasic MN subtypes are tuned to enable their specialized roles in integrating descending commands to coordinate locomotor behavior.

What mechanism(s) might be responsible for the two-fold higher Ca^2+^ influx per action potential at MN-Is release sites over -Ib? While the abundance of Ca_V_2 Ca^2+^ channels at AZs can increase in the context of presynaptic plasticity (Cao & Tsien, 2010; Ghelani et al., 2022; Goel, Bergeron, et al., 2019; Gratz et al., 2019; Sheng et al., 2012), distinctions in Cac abundance do not appear to explain the differences in Ca^2+^ influx at tonic vs phasic release sites. Instead, we consider several possibilities that are not mutually exclusive. First, it is possible that distinct splice variants of *cac* are expressed at MN-Is vs -Ib AZs. Indeed, there are at least 15 predicted splice isoforms of *cac* (FlyBase, 2022), and there is some evidence for differential functions of these isoforms (Lembke et al., 2018). Recent efforts at transcriptomic profiling of the fly larval nervous system and even MN subtypes may shed light on this possibility (FlyAtlas, 2022; Vicidomini et al., 2021). Secondly, another interesting possibility is that there may be different action potential waveforms that invade MN-Is vs -Ib terminals. For example, even relatively moderate differences in the rate of re-polarization can have a large impact on Ca^2+^ influx through voltage-gated Ca^2+^ channels (Schneggenburger, 1996) Voltage imaging of MN-Ib action potentials has been described (Ford & Davis, 2014), but how this compares to action potentials driven in MN-Is is unknown. Thirdly, post-translational modifications of Cac might impart distinct functional states at tonic vs phasic release sites, including RNA editing or changes in the phosphorylation state (Tedford & Zamponi, 2006; Zamponi & Currie, 2013). Gaining mechanistic insight into why phasic release sites experience two-fold increases in Ca^2+^ will be an area for future research to help explain the enhanced release probability of phasic AZs.

### Unique features of AZ architecture and coupling at MN-Is vs -Ib

It would appear counterintuitive, that the “strong” phasic release sites would be represented by less scaffold protein in more compact AZ structures relative to “weak” tonic AZs. AZs represented by a greater presence of scaffold, such as BRP and RBP are typically interpreted as having an enhanced release probability at the fly NMJ (Cunningham et al., 2022; Gratz et al., 2019; Newman et al., 2017) However, our analysis highlights two key aspects of AZ nanostructure that likely contribute to enhanced release at phasic AZs despite their reduced size. First, while BRP and RBP scaffold proteins are indeed reduced at Is vs Ib AZs, Cac, the channel that directly gates Ca^2+^ entry at AZs, is present in similar amounts. Secondly, STED super resolution imaging revealed a major increase in the area ratio of Cac relative to RBP and BRP at Is vs Ib AZs, while the area ratio of RBP to BRP remained similar. A complementary super resolution imaging approach to STED, dSTORM, has also documented a “compaction” of AZ structures at MN-Is relative to Ib (Ehmann et al., 2014)Msrestrani paper – check again they had 1s data, maybe in supplement. This compaction of AZ structure does appear capable of enhancing neurotransmitter release, as a similar compaction has been reported for Ib AZs following a form of adaptive plasticity known as presynaptic homeostatic potentiation (PHP) (Dannhäuser et al., 2022; Goel & Dickman, 2021; Mrestani et al., 2021). Furthermore, an intriguing recent study added an additional layer of regulation of Cac at AZs, where the nanoscale mobility and localization within individual MN-Ib AZs was subject to dynamic regulation at basal and plasticity states (Ghelani et al., 2022). Although these dynamics were not studied at MN-Is AZs, distinctions in Cac mobility and dynamics likely also contribute to the functional differences in tonic vs phasic release sites. While it is clear that smaller, more compact AZ structures can drive enhanced neurotransmitter release, the mechanisms remain unclear.

In addition to enhanced Ca^2+^ influx at MN-Is AZs, there is also an increased vesicle coupling at these phasic release sites compared to MN-Ib. It appears that release at MN-Is is calibrated to be maximal at physiologic Ca^2+^ levels, and the dual levers of Ca^2+^ levels and vesicle coupling may enable a tunable system to permit a tight regulation of transmission when necessary, while still allowing the flexibility necessary to change according to plasticity needs. For example, at lowered Ca^2+^ conditions, PHP selectively targets MN-Ib for enhanced release (Newman et al., 2017), while at elevated Ca^2+^ both tonic and phasic inputs are modulated (Aponte-Santiago & Littleton, 2020; Genç & Davis, 2019; Sauvola et al., 2021). It is tempting to speculate that differential utilization of Unc13 isoforms may contribute to the distinct coupling observed at Is vs Ib AZs, with Unc13A specifying tightly coupled vesicle fusion close to Ca^2+^ channels while loosely coupled vesicles depend on Unc13B and localization farther from Ca^2+^ sources (Reddy-Alla et al., 2017). This work lays a foundation for exciting areas of future research, including how tonic and phasic release sites may be differentially targeted during various forms of plasticity.

## MATERIALS AND METHODS

### Fly stocks

*Drosophila* stocks were raised at 25°C on standard molasses food. The *w^1118^* strain is used as the wild-type control unless otherwise noted as this is the genetic background in which all genotypes are bred. The *UAS-CEPIA1er-P2A-RFP* stock was generated in this study. The genotypes and sources of all stocks used in this study are details in the Key Resources Table.

### Molecular Biology

To generate the *UAS-CEPIAer-hPDGFR-TM-P2A-TagRFP-hPDGFR-TM* transgene (abbreviated to *UAS-CEPIAer-P2A-RFP*), we obtained the sequence of *G-CEPIA1er* (Suzuki et al., 2014) and biosynthesized the cDNA after codon optimization for *Drosophila*. The CEPIA1er sequence was then cloned into a pJFRC14 plasmid backbone containing hPDFGR-TM::P2A::TagRFP::hPDFGR-TM (donated by Gregory T. Macleod lab) using Gibson assembly. The final CEPIA1er::hPDFGR-TM::P2A::TagRFP::hPDFGR-TM construct was transformed into chromosome III of *w^1118^* by Bestgene, Inc (Chino Hills, CA) using PhiC31 integrase-mediated attP site-specific (VK27) transgenesis and subsequently balanced.

### Electrophysiology

All dissections and two-electrode voltage clamp (TEVC) recordings were performed as described (Kikuma et al., 2019) using modified hemolymph-like saline (HL-3) containing: 70mM NaCl, 5mM KCl, 10mM MgCl_2_, 10mM NaHCO_3_, 115mM Sucrose, 5mM Trehelose, 5mM HEPES, and CaCl_2_ of specified concentration, pH 7.2, from cells with an initial resting potential between −60 and −75 mV, and input resistances >6 MΩ. For EGTA sensitivity assays, larval fillets were incubated in 0 Ca^2+^ modified HL-3 supplemented with 25uM EGTA-AM 10 min, then washed with HL-3 three times before recording in standard saline. Recordings were performed on an Olympus BX61 WI microscope using a 40x/0.80 NA water-dipping objective and acquired using an Axoclamp 900A amplifier, Digidata 1440A acquisition system and pClamp 10.5 software (Molecular Devices). Miniature excitatory postsynaptic currents (mEPSCs) were recorded in the absence of any stimulation with a voltage clamp of −80 mV, and low pass filtered at 1 kHz. All recordings were made on abdominal muscle 6, segment A3 of third-instar larvae with the leak current never exceeding 5 nA. mEPSCs were recorded for 60 seconds and analyzed using MiniAnalysis (Synaptosoft) and Excel (Microsoft) software. The average mEPSC amplitude and total charge transfer values for each NMJ were obtained from approximately 100 events in each recording. Excitatory postsynaptic currents (EPSCs) were recorded by delivering 20 electrical stimulations at 0.5 Hz with 0.5 msec duration to motor neurons using an ISO-Flex stimulus isolator (A.M.P.I.) with stimulus intensities set to avoid multiple EPSCs.

The readily releasable pool (RRP) size was estimated as described (Goel, Li, et al., 2019). EPSCs were evoked with a 60 Hz, 60 stimulus train while recording in HL-3 supplemented with 3 mM Ca^2+^. A line fit to the linear phase (stimuli # 18–30) of the cumulative EPSC data was back-extrapolated to time 0. The RRP value was estimated by determining the extrapolated EPSC value at time 0 and dividing this value by the average mEPSP amplitude. Data used in the variance-mean plot was obtained from TEVC recordings using an initial 0.5 mM Ca^2+^ concentration, which was subsequently increased to 1.5, 3.0, and 6.0 mM through saline exchange using a peristaltic pump (Langer Instruments, BT100-2J). EPSC amplitudes were monitored during the exchange, and 30 EPSC recordings (0.5 Hz stimulation rate) were performed in each condition. To obtain the variance-mean plot, the variance (squared standard deviation) and mean (averaged evoked amplitude) were calculated from the 30 EPSCs at each individual Ca^2+^ concentration. The variance was then plotted against the mean for each concentration using MATLAB software (MathWorks, USA). One additional data point, in which variance and mean are both theoretically at 0, was used for Ca^2+^-free saline. Data from these five conditions were fit with a standard parabola (variance=Q*Ī -Ī2/N), where Q is the quantal size, Ī is the mean evoked amplitude (x-axis), and N is the functional number of release sites. N, as a parameter of the standard parabola, was directly calculated for each cell by best parabolic fit.

### Immunocytochemistry

Third-instar larvae were dissected in ice cold 0 Ca^2+^ HL-3 and immunostained as described (Chen and Dickman, 2017; Goel et al., 2017; Li et al., 2021). In brief, larvae were either fixed in 100% ice-cold ethanol for 5 min or 4% paraformaldehyde (PFA) for 10 min. Larvae were then washed with PBS containing 0.1% Triton X-100 (PBST) for 30 min, blocked with 5% Normal Donkey Serum followed by overnight incubation in primary antibodies at 4°C. Preparations were then washed 3x in PBST, incubated in secondary antibodies for 2 hours at room temp, washed 3x in PBST, and equilibrated in 70% glycerol. Prior to imaging, samples were mounted in VectaShield (Vector Laboratories). The guinea pig anti-RBP polyclonal antibody was generated using a synthetic peptide VLSKGKDLFGKF as described (Liu et al., 2011) and affinity purified. Additional details of all antibodies, their source, dilution used, and references are listed in the Key Resources Table.

### Confocal imaging and analysis

Confocal images were acquired with a Nikon A1R Resonant Scanning Confocal microscope equipped with NIS Elements software and a 100x APO 1.4NA oil immersion objective using separate channels with four laser lines (405 nm, 488 nm, 561 nm, and 647 nm) as described (Perry et al., 2017). For fluorescence intensity quantifications of BRP, RBP and CAC, z-stacks were obtained on the same day using identical gain and laser power settings with z-axis spacing of 0.20 μm for all samples within an individual experiment. Raw confocal images were deconvolved with SVI Huygens Essential 22.10 using built-in Express settings. 3D surface rendering images were then generated by Huygens for display. Maximum intensity projections were utilized for quantitative image analysis using the general analysis toolkit of NIS Elements software. Immunofluorescence intensity levels were quantified by applying intensity thresholds and filters to binary layers in the 405 nm, 488 nm, and 561 nm channels. The mean intensity for each channel was quantified by obtaining the average total fluorescence signal for each individual punctum and dividing this value by the puncta area. A mask was created around the HRP channel, used to define the neuronal membrane, and only puncta within this mask were analyzed to eliminate background signals. All measurements based on confocal images were taken from NMJs acquired from at least six different animals.

### STED imaging and analysis

Stimulated Emission Depletion (STED) super resolution microscopy was performed with an Abberior STEDYCON system mounted on a Nikon Eclipse FN1 upright microscope using four excitation lasers (640 nm, 561 nm, 488 nm and 405 nm), a pulsed STED laser of 775 nm, and three avalanche photodiode detectors that operate in a single photon counting mode. Multichannel 2D STED images were acquired using a 100x Nikon Plan APO 1.45NA oil immersion objective with 15 nm fixed pixel size and 10 μsec dwell time using 15x line accumulation in photon counting mode and field of view of 1-2 boutons. 2-3 secondary dyes were used, Abberior STAR Red combined with Alexa Fluor 594 and ATTO 490LS, and all were depleted at 775 nm. For the STED channel, time gating was set at 1 nsec with a width of 6 nsec for all channels. Fluorescence photons were counted sequentially by pixel with the respective avalanche photodiode detector (STAR RED and ATTO490LS: 675 ± 25 nm, Alexa Fluor 594: 600 ± 25 nm, Alexa Fluor 488: 525 ± 25 nm). Raw STED images are deconvolved with the SVI Huygens software using the default settings and theoretical point spread functions for STED microscope. Covered areas of each protein were based on the raw STED images in red and infrared channels and quantified with the general analysis toolkit of NIS Elements software (Version 4.2). Active zones with optimal planar orientation were manually selected and the area and equivalent diameter of each protein was determined by applying an intensity threshold to mask layers in 600 nm and 675 nm channels, and a fixed intensity threshold value was applied to the same channels across samples. The nanomodules of each BRP puncta were quantified with using a bright spot detection algorithm in NIS Elements software with the same threshold setting for each image. The height of each active zone was assessed by measuring the maximum vertical distance between the BRP and CAC signal from manually selected active zones with optimal orientation based on the vertical view of BRP donuts. All measurements were based on 2D STED images that were taken from NMJs of M6 in segments A3 acquired from at least four different animals.

### Live imaging and analysis

#### ex-vivo intracellular Ca^2+^ imaging using rhod-dextran and fura dextran

Female 3^rd^ instar larvae were fillet dissected in chilled Schneider’s insect medium as illustrated previously (Rossano & Macleod, 2007), and the nerve was forward-filled with a 10,000 MW dextran-conjugated Ca^2+^-indicator, rhod dextran (Cat No. R34677, Invitroen), in constant ratio with a 10,000 MW dextran-conjugated Ca^2+^-insensitive fluorophore, AF647 dextran (Cat No. D22914, Invitrogen), as described previously (Chouhan et al., 2010). Ca^2+^ imaging was performed in HL6 (Macleod et al., 2002) with 2 mM Ca^2+^ and 15 mM Mg^2+^ under a water-dipping 100X 1.0NA objective of an Olympus BXWI50 microscope, fitted with an EMCCD camera (Andor Technology, model DU860; South Windsor, CT). Rhod was excited by a DG4 150W Xenon bulb wavelength-switcher (Sutter Instruments) with 543/22 nm light reflected off a 552 nm dichroic mirror (DM); AF647 was similarly excited with 628/40 nm light off a 660 nm DM. Emission was collected through a Sutter instruments lambda 10-B filter wheel (rhod, 593/40 nm; AF647, 692/40 nm.). Filters and dichroic mirrors were obtained from Chroma Technology (Bellows Falls, VT, USA) or Semrock (Lake Forest, IL, USA). Several images of AF647 were captured before nerve stimulation, but rhod images were captured before, during, and after stimulation, at 100 frames-per-second.

Intracellular Ca^2+^ concentration ([Ca^2+^]_i_), and the number of Ca^2+^ ions entering through a single AZ (on average) were determined as described by Lu and others (2016). Briefly, the first step was to convert the ratio of rhod to AF647 fluorescence intensity to [Ca^2+^]_i_ using Equation 5 of (Grynkiewicz et al., 1985). Values of R_max_ were obtained *in situ* through incubation of preparations in HL6 containing 10mM Ca^2+^ and 100 uM ionomycin (Cat. No. I9657; Sigma) and R_min_ values were obtained by incubating preparations in Ca^2+^-free HL6 with 1 mM EGTA and 100 uM BAPTA-AM (Cat No. B6769; Invitrogen) (1% DMSO) for 20 min. The K_D_ value used for rhod-dextran was the batch specific value of 3 uM. The second step was to calculate the total change in [Ca^2+^] in each terminal (both bound and unbound by rhod and the endogenous Ca^2+^ buffers) using Equation 4 of (Helmchen et al., 1996). Here, we made two simplifying assumptions; that the endogenous Ca^2+^ binding ratio (K_S_) was the same between terminals (48.35) as concluded by (Justs et al., 2022), and that the Ca^2+^ extrusion rate (gamma) was no different between terminals (987 nM s^-1^); the latter assumption being consistent with similar rates of decay (tau) in the two terminals types when the exogenous dye concentration is shown to be no different between terminals types (Lu et al., 2016). The third step was to calculate the total number of Ca^2+^ ions that entered each terminal using equation 4 of Lu et al (2016), where the total change in [Ca^2+^]_i_ concentration is multiplied by the average volume of each terminal on muscle 6 (Ib: 310 ± 49 um^3^; Is: 90 ± 49 um^3^; N=6; Lu et al., 2016) and Avagadro’s constant. The final step was to calculate the number of Ca^2+^ ions that entered through each AZ on each terminal on muscle 6 by dividing by the number of AZs for each terminal established through 3D EM reconstruction (Ib: 747 ± 114; Is: 223 ± 66; N=6; Lu et al., 2016).

For fura imaging, a similar protocol was used as for rhod-dextran imaging above, previously described by (Chouhan et al., 2010). Briefly, nerve terminals were forward-filled with fura-dextran (Cat No. F3029 Invitrogen; K_D_=865 nM) and imaged in HL6 with 2 mM Ca^2+^ on the same “rig” as described above, but using different filters and at a slower acquisition rate 4 images/s, yielding 2 ratios/s). Fura was alternately excited by 340/26 and 387/11 nm light reflected off a 409 nm DM, with emitted light collected through a 510/84 nm filter. Fluorescence signals were converted to [Ca^2+^]_i_ using Equation 5 of (Grynkiewicz et al., 1985) after determining values of R_max_ and R_min_, as above.

#### ex-vivo intracellular Ca^2+^ imaging using GCaMP6m and ex-vivo intracellular pH imaging using Superecliptic-pHluorin

Fillet dissections of female larvae were performed in chilled Schneider’s insect medium as above, before transferring into HL6 with CaCl_2_ added to 0.1 mM. Although brain hemispheres were removed, hemisegment nerves were not severed. Images were collected using a Nikon 60X, 1.20 NA, Plan Apochromat VC water-immersion objective on a Nikon A1R confocal microscope fitted with GaAsP detectors. Resting Ca^2+^ levels and pH were compared between terminals by estimating the fluorescence intensity ratio of GCaMP6m (Ca^2+^) to either mCherry or myristoylated FusionRed, and SE-pHluorin (pH) to mCherry. The fluorophores were scanned sequentially using 488 nm ex. / 520 nm em., then 561 nm ex. / 590 nm em with a pinhole of 3 Airy units. Images were analyzed on ImageJ and mean intensity values of fluorescence from ROIs surrounding boutons and R0Is of bouton backgrounds were exported to Microsoft Excel. In each case, ratios represent the green fluorescence of the Ca^2+^ or pH-sensitive fluorophore divided by the red fluorescence of the insensitive fluorophores (bouton-background in each case).

#### in vivo and ex-vivo extracellular Ca^2+^ imaging using CEPIA1er

The extracellular Ca^2+^ concentration ([Ca^2+^]_e_) calibration curve of CEPIA1er fluorescence relative to RFP fluorescence was obtained using a serial dilution of chilled 10 mM [Ca^2+^] HL6 to generate a [Ca^2+^]_e_ series of 0.1 mM, 0.5 mM, 1 mM, 3 mM, 5 mM and 10 mM; each with 20 mM [Mg^2+^] and pH 7.2. Female 3^rd^ instar larvae expressing CEPIA1er-P2A-RFP pan-neuronally (nSyb-Gal4) were fillet dissected in chilled Schneider’s solution. The ventral ganglion was removed, and the preparation was flipped (ventral side down), to mimic *in vivo* imaging conditions where images were captured through the cuticle of larvae. A groove was excavated in the Sylgard base of the chamber to allow adequate exchange of saline. Images were collected from type-Ib terminals using the confocal microscope as described for GCaMP and its red fluorophore counterparts but with a number of protocol modification. CEPIA1er to RFP ratio estimates were collected for all 6 Ca^2+^ concentrations from each of 7 *ex vivo* preparations; 4 series were collected in an increasing series of Ca^2+^ concentrations, and 3 were collected in a decreasing series. When exchanging saline, the preparation was rinsed twice with the new [Ca^2+^]_e_ saline, then once more prior to imaging when it was covered with a glass cover slip. All *ex vivo* images were taken from the segment exposed to the Sylgard groove. Undissected larvae were prepared for imaging by constraining them in a snug form-fitting recess in a Sylgard tablet and covered with a glass cover-slip leaving only the posterior spiracles open to the air. MN6/7-Ib boutons with minimal background fluorescence were selected for *in vivo* imaging. Boutons were scanned first with 561 nm light (RFP; 590 nm em.), and then with 488 nm light (CEPIA1er; 520 nm em.). No bleed-through could be detected between channels. Images were analyzed on ImageJ, as described above, and the ratio represents the CEPIA signal divided by the RFP signal. When a series of 6 images had to be collected during ex vivo calibration measurements, RFP bleached more rapidly than CEIPIA1er. To control for this effect, and to allow for a direct comparison between *ex vivo* calibrations and the *in vivo* estimates where only one image was captured, RFP fluorescence captured in the first image was used to calculate the CEPIA1er to RFP ratio for each *ex vivo* calibration series. Lastly a fit of the calibration curve was performed in MATLAB using the Hill equation, and the *in vivo* hemolymph [Ca^2+^] was obtained by finding the intercept of the average *in vivo* ratio with *ex vivo* [Ca^2+^]_e_ calibration curve.

### Statistical analysis

Data were analyzed using GraphPad Prism (version 8.0), MiniAnalysis (Synaptosoft), SVI Hugyens Essential (Version 22.10), or Microsoft Excel software (version 16.22). Sample values were tested for normality using the D’Agostino & Pearson omnibus normality test which determined that the assumption of normality of the sample distribution was not violated. Data were then compared using either a one-way ANOVA and tested for significance using a Tukey’s multiple comparison test or using an unpaired 2-tailed Student’s t-test with Welch’s correction. In all figures, error bars indicate ±SEM, with the following statistical significance: p<0.05 (*), p<0.01 (**), p<0.001 (***), p<0.0001 (****); ns=not significant. Additional statistics and sample number values (n) for all experiments are summarized in Supplemental Table S1.

## Supporting information

Key Resources Table

## ACKNOWLEDGEMENTS

We thank Kris Fertig, Joachim Fischer, and John Waka from Abberior Instruments for technical advice on STED microscopy and Bryan Stewart (U of Toronto, Canada) for important discussions. We acknowledge the Developmental Studies Hybridoma Bank (Iowa, USA) for antibodies used in this study and the Bloomington Drosophila Stock Center for fly stocks (NIH P40OD018537). This work was supported by grants from the National Institutes of Health (NS103906) to G.M. and (NS126654 and NS111414) to D.D.

## AUTHOR CONTRIBUTIONS

K.H., G.M., and D.D. designed the research project, K.H., Y.H., X.L., R.H., D.R., G.M., T.F., K.J., O.M., and S.P. performed experiments and analyzed the data. The manuscript was written by K.H., G.M., and D.D. with feedback from the other authors.

---

**Figure S1:**
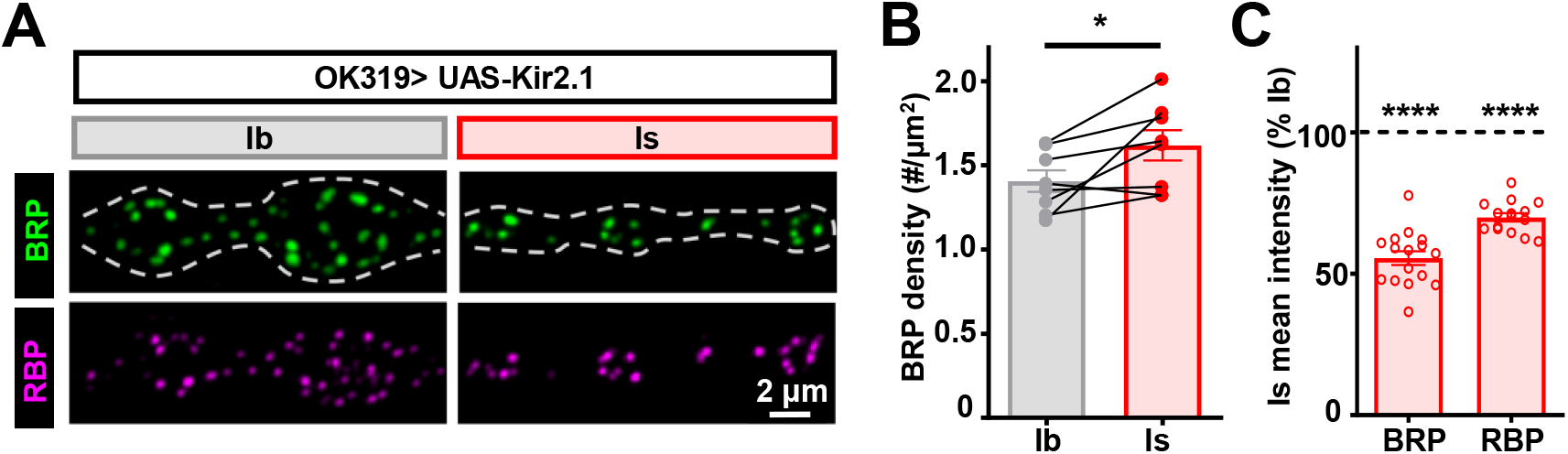
Tonic vs phasic firing patterns do not determine AZ distinctions. **(A)** Tonic vs phasic firing patterns are disrupted by expression of the inwardly rectifying potassium channel *Kir2.1* in both MN-Is and -Ib (OK319>Kir2.1: *w;OK319/+;UAS-Kir2.1-GFP/+*). Representative confocal images of muscle 6 NMJs in these larvae immunostained with antibodies against the active zone components BRP and RBP at MN-Ib and MN-Is. The dashed line represents the neuronal membrane indicated by the anti-HRP signal. **(B-C)** Similar BRP puncta density and mean intensities were observed compared to wild type (Fig. 6) despite altered activity patterns, suggesting that tonic vs phasic activity is not required for the dimmed active zone components characteristic of MN-Is. Error bars indicate ± SEM. ****p<0.0001 *p<0.05. Additional statistical details are shown in Supplemental Table S1.

## Notes

### Competing Interest Statement

The authors have declared no competing interest.

