## Supplementary Table 1 for "Physiologic and nanoscale distinctions define glutamatergic synapses in tonic vs phasic neurons"

**Supplementary Table 1: Absolute values for normalized data and additional statistical details.** The figure and panel, genotype, extracellular calcium concentration and conditions are noted. Average values (with standard error of the mean noted in parentheses), data samples (n), and statistical significance tests are shown for all data.

| **Figure** | **Label** | **[Ca^2+^] (mM)** | **Genotype** | **mEPSP amplitude (mV)** | **EPSC amplitude (nA)** | **QC** | **mEPSP frequency (Hz)** | **Rinput (MΩ)** | **Resting potential (mV)** | **n** | **P Value (significance): mEPSP, EPSC, QC, mEPSP freq.** |
| --- | --- | --- | --- | --- | --- | --- | --- | --- | --- | --- | --- |
| 1A, D, E, F | wild type | 0.4 | *w^1118^* | 1.00 (±0.03) | 42.86 (±4.14) | 41.05 (±2.68) | 3.35 (±0.25) | 12.75 (±0.79) | -68.2 (±2.28) | 12 | - |
| 1A, D, E, F | Ib only (Is> BoNT) | 0.4 | *w*; +; *R27E09-GAL4*/*UAS-BoNT-C* | 0.72 (±0.04) | 10.02  (±1.70) | 14.62 (±2.95) | 2.35 (±0.14) | 12.44 (±1.14) | -65.79 (±2.60) | 9 | <0.0001 (****), <0.0001 (****),  <0.0001 (****), <0.05 (*) |
| 1A, D, E, F | Is only (Ib>BoNT) | 0.4 | *w*; +; *dHb9-GAL4*/*UAS-BoNT-C* | 1.19 (±0.05) | 29.10 (±2.96) | 24.72 (±2.78) | 2.04 (±0.31) | 13.60 (±0.69) | -65.13 (±2.02) | 10 | <0.01 (**), <0.01 (**),  <0.0001 (****), <0.01 (**) |
| 1E, F | wild type | 1.0 | *w^1118^* | 1.05 (±0.08) | 108.72 (±4.50) | 107.53 (±1.00) | 3.01 (±0.23) | 5.25 (±0.52) | -68.16 (±1.52) | 8 | - |
| 1E, F | Ib only | 1.0 | *w*; +; *R27E09-GAL4*/*UAS-BoNT-C* | 0.77 (±0.05) | 32.14 (±2.78) | 42.42 (±3.37) | 2.44 (±0.10) | 6.00 (±0.25) | 68.60 (±1.31) | 9 | <0.01 (**), <0.0001 (****), <0.0001 (****), <0.05 (*), |
| 1E, F | Is only | 1.0 | *w*; +; *dHb9-GAL4*/*UAS-BoNT-C* | 1.15 (±0.05) | 90.74 (±2.78) | 79.03 (±2.21) | 1.52 (±0.05) | 10.38 (±0.40) | 67.53 (±1.54) | 8 | 0.42 (ns), <0.01 (**), <0.01 (**), <0.0001 (****) |
| 1B, E, F | wild type | 1.8 | *w^1118^* | 1.06 (±0.04) | 185.93 (±11.17) | 178.20 (±12.23) | 3.19 (±0.47) | 11.00 (±0.89) | -64.88 (±1.87) | 12 | - |
| 1B, E, F | Ib only | 1.8 | *w*; +; *R27E09-GAL4*/*UAS-BoNT-C* | 0.86 (±0.07) | 59.9 (±3.35) | 74.35 (±13.38) | 0.61 (±0.08) | 11.40 (±1.08) | -66.04 (±2.57) | 6 | <0.05 (*), <0.0001 (****), <0.0001 (****), <0.01 (**) |
| 1B, E, F | Is only | 1.8 | *w*; +; *dHb9-GAL4*/*UAS-BoNT-C* | 1.29 (±0.06) | 131.54 (±10.11) | 102.8 (±7.47) | 1.20 (±0.17) | 9.80 (±0.86) | -69.64 (±2.70) | 6 | <0.01 (**), <0.01 (**), <0.001 (***), <0.01 (**) |
| 1C, E, F | wild type | 3.0 | *w^1118^* | 1.07 (±0.03) | 208.75 (±11.83) | 197.91 (±11.09) | 3.11 (±0.37) | 10.10 (±0.76) | -79.69 (±1.80) | 20 | - |
| 1C, E, F | Ib only | 3.0 | *w*; +; *R27E09-GAL4*/*UAS-BoNT-C* | 0.71 (±0.03) | 107.79 (±7.38) | 149.57 (±12.55) | 1.95 (±0.25) | 5.60 (±0.27) | -70.4 (±1.85) | 10 | <0.0001 (****), <0.0001 (****), <0.01 (**), <0.01 (**) |
| 1C, E, F | Is only | 3.0 | *w*; +; *dHb9-GAL4*/*UAS-BoNT-C* | 1.31 (±0.06) | 122.9 (±3.59) | 99.96 (±5.30) | 1.54 (±0.17) | 7.80 (±1.09) | -71.25 (±1.36) | 11 | <0.001 (***), <0.0001 (****), <0.0001 (****), <0.001 (***) |
| 1E, F | wild type | 6.0 | *w^1118^* | 1.06 (±0.07) | 239.71 (±10.3) | 229.88 (±11.20) | 3.02 (±0.18) | 6.00 (±0.20) | -68.16 (±1.53) | 8 | - |
| 1E, F | Ib only | 6.0 | *w*; +; *R27E09-GAL4*/*UAS-BoNT-C* | 0.74 (±0.03) | 119.05 (±9.20) | 161.76 (±13.57) | 2.11 (±0.15) | 5.1 (±0.46) | -72.14 (±0.84) | 10 | <0.001 (***), <0.0001 (****), <0.001 (***), <0.01 (**) |
| 1E, F | Is only | 6.0 | *w*; +; *dHb9-GAL4*/*UAS-BoNT-C* | 1.27 (±0.06) | 138.24 (±5.18) | 109.84 (±3.67) | 1.49 (±0.23) | 4.75 (±0.39) | -71.82 (±0.91) | 8 | <0.05 (*), <0.0001 (****), <0.0001 (****), <0.0001 (****) |

| **Figure** | **Label** | **[Ca^2+^] (mM)** | **Genotype** | **%EPSC2/EPSC1** | **Rinput (MΩ)** | **Resting potential (mV)** | **n** | **P Value (significance): EPSC2/EPSC1** |
| --- | --- | --- | --- | --- | --- | --- | --- | --- |
| 2A, B | Ib only | 0.4 | *w*; +; *R27E09-GAL4*/*UAS-BoNT-C* | 241.70 (±22.83) | 12.44 (±1.14) | -65.79 (±2.60) | 9 | - |
| 2A, B | Is only | 0.4 | *w*; +; *dHb9-GAL4*/*UAS-BoNT-C* | 118.78 (±9.66) | 13.60 (±0.69) | -65.13 (±2.02) | 10 | <0.0001 (****) |
| 2A, B | Ib only | 1.8 | *w*; +; *R27E09-GAL4*/*UAS-BoNT-C* | 133.76 (±3.45) | 11.40 (±1.08) | -66.04 (±2.57) | 6 | - |
| 2A, B | Is only | 1.8 | *w*; +; *dHb9-GAL4*/*UAS-BoNT-C* | 74.89 (±2.67) | 9.80 (±0.86) | -69.64 (±2.70) | 6 | <0.0001 (****) |
| 2A, B | Ib only | 3.0 | *w*; +; *R27E09-GAL4*/*UAS-BoNT-C* | 120.00 (±5.19) | 1.95 (±0.25) | -70.4 (±1.85) | 10 | - |
| 2A, B | Is only | 3.0 | *w*; +; *dHb9-GAL4*/*UAS-BoNT-C* | 85.24 (±2.46) | 1.54 (±0.17) | -71.25 (±1.36) | 11 | <0.0001 (****) |
| 2A, B | Ib only | 6.0 | *w*; +; *R27E09-GAL4*/*UAS-BoNT-C* | 81.70 (±3.79) | 11.20 (±0.54) | -73.03 (±3.23) | 10 | - |
| 2A, B | Is only | 6.0 | *w*; +; *dHb9-GAL4*/*UAS-BoNT-C* | 73.20 (±2.49) | 9.00 (±0.61) | -63.79 (±3.04) | 8 | 0.08 (ns) |

| **Figure** | **Label** | **[Ca^2+^] (mM)** | **Genotype** | **Failure rate (%)** | **Rinput (MΩ)** | **Resting potential (mV)** | **n** | **P Value (significance): Failure rate** |
| --- | --- | --- | --- | --- | --- | --- | --- | --- |
| 3A, B | Ib only | 0.1 | *w*; +; *R27E09-GAL4*/*UAS-BoNT-C* | 95 (±1.58) | 6.28 (±0.58) | -63.73 (±2.90) | 11 | <0.0001 (****) |
| 3A, B | Is only | 0.1 | *w*; +; *dHb9-GAL4*/*UAS-BoNT-C* | 65 (±5.32) | 6.71 (±0.61) | -58.57(±1.70) | 7 | - |

| **Figure** | **Label** | **[Ca^2+^] (mM)** | **Genotype** | **Cumulative EPSC (nA)** | **Estimated RRP size** | **Rinput (MΩ)** | **Resting potential (mV)** | **n** | **P Value (significance): Cum. EPSC, RRP size** |
| --- | --- | --- | --- | --- | --- | --- | --- | --- | --- |
| 2C, D | Ib only | 3.0 | *w*; +; *R27E09-GAL4*/*UAS-BoNT-C* | 567.35 (±59.70) | 817.41 (±93.72) | 5.86 (±0.50) | -63.76 (±1.27) | 10 | - |
| 2C, D | Is only | 3.0 | *w*; +; *dHb9-GAL4*/*UAS-BoNT-C* | 493.28 (±56.58) | 358.29 (±37.25) | 5.86 (±0.28) | -72.7 (±3.18) | 7 | 0.37 (ns), <0.001 (***) |
| 3C, D | Ib only | 6.0 | *w*; +; *R27E09-GAL4*/*UAS-BoNT-C* | 514.40 (±19.32) | 703.21 (±42.53) | 11.20 (±0.54) | 73.03 (±3.23) | 10 | - |
| 3C, D | Is only | 6.0 | *w*; +; *dHb9-GAL4*/*UAS-BoNT-C* | 538.75 (±36.72) | 427.64 (±29.35) | 9.00 (±0.61) | 63.79 (±3.04) | 8 | 0.52 (ns), <0.0001 (****) |

| **Figure** | **Label** | **Genotype** | **Release site #** | **Rinput (MΩ)** | **Resting potential (mV)** | **n** | **P Value (significance): Release site #** |
| --- | --- | --- | --- | --- | --- | --- | --- |
| 4C, D | Ib only | *w*; +; *R27E09-GAL4*/*UAS-BoNT-C* | 1222 (±213) | 5.80 (±0.74) | -59.60(±4.28) | 5 | - |
| 4C, D | Is only | *w*; +; *dHb9-GAL4*/*UAS-BoNT-C* | 401 (±52) | 6.00 (±0.63) | -56.8 (±3.01) | 6 | 0.37 (ns), <0.01 (**) |

| **Figure** | **Label** | **[Ca^2+^] (mM)** | **Genotype** | **EGTA-AM** | **EPSC (nA)** | **% of baseline (-EGTA)** | **Rinput (MΩ)** | **Resting potential (mV)** | **n** | **P Value (significance): % baseline (-EGTA)** |
| --- | --- | --- | --- | --- | --- | --- | --- | --- | --- | --- |
| 8D, E | Ib only | 1.8 | *w*; +; *R27E09-GAL4*/*UAS-BoNT-C* | - | -83.98 (±14.43) | - | 5.75 (±0.73) | -58.75 (±3.60) | 4 | - |
| 8D, E | Is only | 1.8 | *w*; +; *dHb9-GAL4*/*UAS-BoNT-C* | - | -130.03 (±14.3) | - | 6.25 (±0.72) | -55.50 (±4.07) | 4 | - |
| 8D, E | Ib only | 1.8 | *w*; +; *R27E09-GAL4*/*UAS-BoNT-C* | + | -46.09 (±8.85) | 54.64 (±4.65) | 5.25 (±0.55) | -58.50 (±3.48) | 4 | - |
| 8D, E | Is only | 1.8 | *w*; +; *dHb9-GAL4*/*UAS-BoNT-C* | + | -95.89 (±10.24) | 73.83 (±1.06) | 5.00 (±0.47) | -52.25 (±3.75) | 4 | <0.01 (**) |

| **Figure** | **Label** | **Genotype** | **Brp puncta #/M6 NMJ** | **n** | **P Value (significance): Brp puncta #/M6 NMJ** |
| --- | --- | --- | --- | --- | --- |
| 4A, B | Ib | *w^1118^* | 198.11 (±10.72) | 9 | - |
| 4A, B | Is | *w^1118^* | 89.89 (±11.5) | 9 | <0.0001 (****) |

| **Figure** | **Label** | **Genotype** | **Indicators (Y axis title)** | **Ratio of indicators** | **n** | **P Value (significance): ratio of indicators** |
| --- | --- | --- | --- | --- | --- | --- |
| 5A | Ib | *w; +; nSyb-GAL4>GCaMP6m-mCherry* | GCaMP6m/mCherry | 1.04 (±0.03) | 11 | - |
| 5A | Is | *w; +; nSyb-GAL4>GCaMP6m-mCherry* | GCaMP6m/mCherry | 0.83 (±0.02) | 11 | <0.0001 (****) |
| 5A | Ib | *w; OK6-GAL4/+; myrFusionRed-GCaMP6m/+* | GCaMP6m/myr-FusionRed | 1.19 (±0.06) | 12 | - |
| 5A | Is | *w; OK6-GAL4/+; myrFusionRed-GCaMP6m/+* | GCaMP6m/myr-FusionRed | 0.86 (±0.05) | 12 | <0.001 (***) |
| 5B | Ib | *w; +; nSyb-GAL4>SE-pHluorin-mCherry* | SE-pHluorin/mCherry | 1.97 (±0.01) | 12 | - |
| 5B | Is | *w; +; nSyb-GAL4>SE-pHluorin-mCherry* | SE-pHluorin/mCherry | 1.76 (±0.02) | 12 | <0.0001 (****) |

| **Figure** | **Label** | **Ca^2+^ Dye** | **Genotype** | **Δ[Ca^2+^]_i_ (nM)** | **Ca^2+^ entry/AZ (10^3^ ions)** | **n** | **P Value (significance): Δ[Ca^2+^]_i_, Ca^2+^ entry/AZ** |
| --- | --- | --- | --- | --- | --- | --- | --- |
| 5D, E, F | Ib | rhod | *w^1118^* | -134.6 (±18.9) | 3.40 (±0.44) | 6 | - |
| 5D, E, F | Is | rhod | *w^1118^* | - 316.5 (±73.8) | 6.79 (±1.31) | 6 | <0.05 (*), <0.05 (*) |

| **Figure** | **Label** | **Ca^2+^ Dye** | **Genotype** | **Stimulation frequency (Hz)** | **[Ca^2+^]_i_ (μM)** | **n** | **P Value (significance): [Ca^2+^]_i_** |
| --- | --- | --- | --- | --- | --- | --- | --- |
| 5I | Ib | fura | *w^1118^* | 0 | 0.057 (±0.009) | 22 | - |
| 5I | Is | fura | *w^1118^* | 0 | 0.057 (±0.010) | 14 | 0.97 (ns) |
| 5I | Ib | fura | *w^1118^* | 20 | 0.303 (±0.025) | 15 | - |
| 5I | Is | fura | *w^1118^* | 20 | 0.492 (±0.052) | 7 | <0.05 (*) |
| 5I | Ib | fura | *w^1118^* | 40 | 0.558 (±0.048) | 15 | - |
| 5I | Is | fura | *w^1118^* | 40 | 0.787 (±0.046) | 11 | <0.05 (*) |
| 5I | Ib | fura | *w^1118^* | 80 | 0.834 (±0.053) | 16 | - |
| 5I | Is | fura | *w^1118^* | 80 | 1.221 (±0.075) | 7 | <0.05 (*) |

| **Figure** | **Label** | **Genotype** | **BRP density (#/μm2)** | **BRP mean intensity** | **RBP mean intensity** | **Cac mean intensity** | **n** | **P Value (significance): BRP density, mean intensity of BRP, RBP, Cac** |
| --- | --- | --- | --- | --- | --- | --- | --- | --- |
| 6B, C | Ib | *Cac^sfGFP-N^; +; +* | 1.33 (±0.04) | 758.25 (±29.65) | 752.33 (±11.71) | 426.42 (±12.68) | 12 | - |
| 6B, C | Is | *Cac^sfGFP-N^; +; +* | 1.51 (±0.04) | 571.67 (±23.78) | 584.58 (±18.38) | 414.67 (±25.01) | 12 | <0.01 (**), <0.0001 (****), <0.0001 (****), 0.57 (ns) |
| S1B, C | Ib | *w; OK319-GAL4> UAS-Kir2.1-GFP; +* | 1.41 (±0.07) | 4791.27 (±395.34) | 2610.71 (±146.31) | - | 13 | - |
| S1B, C | Is | *w; OK319-GAL4> UAS-Kir2.1-GFP; +* | 1.62 (±0.10) | 2184.61 (±88.00) | 1529.18 (±33.20) | - | 13 | <0.05 (*), <0.0001 (****), <0.0001 (****) |

| **Figure** | **Label** | **Genotype** | **BRP ring area (μm2)** | **RBP/BRP area ratio (%)** | **Nanomodules/ring** | **n** | **P Value (significance): BRP ring area (μm^2^), RBP/BRP area ratio (%), nanomodules/ring,** |
| --- | --- | --- | --- | --- | --- | --- | --- |
| 7B, D | Ib | *w^1118^* | 0.0736 (±0.0028) | 60.00 (±3.59) | 4.20 (±0.17) | 22 | - |
| 7B, D | Is | *w^1118^* | 0.0491 (±0.0039) | 62.30 (±4.57) | 2.75 (±0.20) | 13 | <0.0001 (****), 0.69 (ns), <0.0001 (****) |

| **Figure** | **Label** | **Genotype** | **BRP ring diameter (nm)** | **BRP ring height (nm)** | **Cac/BRP area ratio (%)** | **n** | **P Value (significance): BRP ring diameter, height, Cac/BRP area ratio (%)** |
| --- | --- | --- | --- | --- | --- | --- | --- |
| 7D, F | Ib | *Cac^sfGFP-N^; +; +* | 293.88 (±4.50) | 180.05 (±5.64) | 36.33 (±2.27) | 21 | - |
| 7D, F | Is | *Cac^sfGFP-N^; +; +* | 266.94 (±5.26) | 158.99 (±5.54) | 73.40 (±5.64) | 10 | <0.001 (***), <0.05 (*), <0.0001 (****) |

| **Figure** | **[Ca^2+^] (mM)** | **Genotype** | **CEPIAer/RFP** | **n** |
| --- | --- | --- | --- | --- |
| 8B | 0.1 | *w; +; nSyb-GAL4> CEPIAer::PDGFR-P2A-RFP::PDGFR* | 1.55 (±0.30) | 13 |
| 8B | 0.5 | *w; +; nSyb-GAL4> CEPIAer::PDGFR-P2A-RFP::PDGFR* | 1.98 (±0.41) | 13 |
| 8B | 1.0 | *w; +; nSyb-GAL4> CEPIAer::PDGFR-P2A-RFP::PDGFR* | 2.29 (±0.46) | 13 |
| 8B | 3.0 | *w; +; nSyb-GAL4> CEPIAer::PDGFR-P2A-RFP::PDGFR* | 2.58 (±0.51) | 13 |
| 8B | 5.0 | *w; +; nSyb-GAL4> CEPIAer::PDGFR-P2A-RFP::PDGFR* | 2.84 (±0.40) | 13 |
| 8B | 10.0 | *w; +; nSyb-GAL4> CEPIAer::PDGFR-P2A-RFP::PDGFR* | 2.86 (±0.74) | 13 |
| 8B | *in vivo* | *w; +; nSyb-GAL4> CEPIAer::PDGFR-P2A-RFP::PDGFR* | 2.56 (±0.37) | 6 |

| **Figure 8** | **[Mg^2+^] (mM)** | **[Ca^2+^] (mM)** | **Genotype** | **EPSC (nA)** | **Rinput (MΩ)** | **Resting potential (mV)** | **n** | **P Value (significance): EPSC** |
| --- | --- | --- | --- | --- | --- | --- | --- | --- |
| extend | 10 | 1.8 | *w^1118^* | -252 (±7.53) | 5.28 (±0.45) | -55.43 (±3.12) | 7 | - |
| extend | 15 | 1.8 | *w^1118^* | -250 (±11.16) | 5.86 (±0.55) | -57.57(±2.12) | 7 | 0.90 (ns) |
