## Supplementary material for "Physiologic and nanoscale distinctions define glutamatergic synapses in tonic vs phasic neurons": Key Resources Table

**KEY RESOURCE TABLE**

| **REAGENT/RESOURCE** |  | **SOURCE** | **IDENTIFIER** |
| --- | --- | --- | --- |
| **Antibodies** | **Dilution** |  |  |
| Mouse anti-BRP (nc82) | 1:250 | Developmental Studies Hybridoma Bank (DSHB) | AB_2314866 |
| Guinea pig anti-RBP | 1:2000 | This study |  |
| Chicken anti-GFP | 1:400 | Aves Labs | GFP-1020 |
| Alexa Fluor 488 conjugated Goat anti-Horseradish Peroxidase | 1:400 | Jackson ImmunoReserach Laboratories (Jackson) | 123-545-021 |
| Alexa Fluor 488 conjugated secondary antibodies | 1:400 | Jackson | 715-545-150 |
| Cy3-conjugated secondary antibodies | 1:400 | Jackson | 106-165-003 |
| Alexa Fluor 594 conjugated secondary antibodies | 1:400 | Jackson | 106-585-003  103-585-155 |
| Alexa Fluor 647 conjugated Goat anti-Horseradish Peroxidase | 1:400 | Jackson | 123-605-021 |
| ATTO490LS conjugated secondary antibodies | 1:200 | Hypermol | 2109-1MG |
| STAR RED conjugated secondary antibodies | 1:200 | Abberior | STRED-1001  STRED-1006 |
| **Drosophila Strains** |  |  |  |
| UAS-CEPIA1er-hPDGFR-TM-P2A-RFP-hPDGFR-TM |  | This Study |  |
| UAS-myrFusionRed-GCaMP6m |  | (Stawarski et al., 2020) |  |
| UAS-GCaMP6m-mCherry |  | (Ugur et al., 2017) |  |
| UAS-SE-pHluorin-mCherry |  | (Rossano et al., 2017) |  |
| UAS-BoNT-C |  | (Han et al., 2022) |  |
| OK319-GAL4 |  | (Beck et al., 2012) |  |
| *Cac^sfGFP-N^* |  | (Gratz et al., 2019) |  |
| UAS-Kir2.1-GFP |  | Bloomington Drosophila Stock Center (BDSC) | 6596 |
| nSyb-GAL4 |  | BDSC | 51635 |
| OK6-GAL4 |  | BDSC | 64199 |
| dHb9-GAL4 (Ib-Gal4) |  | BDSC | 83004 |
| GMR27E09-GAL4 (Is-Gal4) |  | BDSC | 49227 |
| *w^1118^* |  | BDSC | 5905 |
| **Software and Algorithms** |  |  |  |
| NIS Elements software |  | Nikon | 4.51.01 |
| Huygens Essential |  | SVI | 22.10 |
| Axon pCLAMP |  | Molecular Devices | 10.7 |
| MiniAnalysis |  | Synaptosoft | 6.0.3 |
| GraphPad Prism |  | GraphPad | 8.0.1 |
